# Structures of the 26S proteasome in complex with the Hsp70 cochaperone Bag1 reveal a novel mechanism of ubiquitin-independent proteasomal degradation

**DOI:** 10.1101/2025.01.22.633148

**Authors:** Moisés Maestro-López, Tat Cheung Cheng, Jimena Muntaner, Margarita Menéndez, Melissa Alonso, Andreas Schweitzer, Jorge Cuéllar, José María Valpuesta, Eri Sakata

**Author notes:** These authors contributed equally (Moisés Maestro-López, Tat Cheung Cheng, Jimena Muntaner). **Correspondence authors:** Jorge Cuéllar, José María Valpuesta, Eri Sakata.

## Abstract

The 26S proteasome primarily degrades proteins marked by polyubiquitin chains. Although ubiquitin-independent pathways for proteasomal degradation exist, the mechanisms involved remain poorly understood. Bag1 links the Hsp70 chaperone to the 26S proteasome, recruiting Hsp70-bound aberrant proteins for degradation. Here, we present high-resolution cryo-EM structures of the Bag1-bound 26S proteasome, which reveal unprecedented conformational changes within the 19S regulatory particle. Bag1 binding to the Rpn1 subunit induces a dramatic reconfiguration of AAA+ ATPase subunits, disrupting the canonical spiral staircase conformation and remodeling the central channel architecture. This creates a large cavity above the substrate entry gate of the 20S core particle, enabling the direct entry of Hsp70 clients into the proteolytic chamber. Thus, in this ubiquitin-independent degradation pathway, unfolded proteins can bypass the need for both client ubiquitination and ATP hydrolysis for degradation.

## Introduction

The coordination between the chaperone network and protein degradation pathways is crucial for maintaining cellular proteostasis^1^. While chaperones attempt to refold nascent proteins and abnormal proteins and solubilize aggregated proteins, if unsuccessful, the aberrant proteins are eliminated through regulated degradation^2,3^. In the ubiquitin-proteasome system (UPS), covalent conjugation of polyubiquitin chains on substrates guides specific protein degradation by the 26S proteasome^4,5^. The degradation process starts with the recognition of ubiquitinated substrates by the 19S regulatory particle (RP), followed by their unfolding and translocation through a central channel into the proteolytic chamber of the 20S core particle (CP)^5,6^. Six AAA+ ATPase subunits, Rpt1-6, unfold substrates by processive threading through the central channel, fueled by sequential ATP hydrolysis^7–9^. Proteasome function is finely tuned by transiently associated cofactors, as represented by ubiquitin-like (UBL) and ubiquitin-associated (UBA) proteins (Rad23, Dsk2/Ubqln), which assist with recruiting polyubiquitinated substrates to the 26S proteasome^10–12^. The UBL domain can be found in various proteasome cofactors with different functions, including the deubiquitinating enzyme (DUB) Ubp6/Usp14^13^, the phosphatase Ublcp1^14^, and the Hsp70 cochaperone Bag1^15,16^.

Besides the degradation of ubiquitinated proteins, the proteasome can also process certain proteins through ubiquitin (Ub)-independent pathways. For example, unfolded proteins^17,18^, intrinsically disordered proteins (IDPs)^19,20^, and some specific regulatory proteins which require rapid turnover during signal and stress responses^21–24^, have been reported to be eliminated through ubiquitin-independent mechanisms, sometimes with the assistance of adaptor proteins. However, the precise mechanisms by which the 26S proteasome executes ubiquitin-independent degradation remains poorly understood.

Hsp70s are a large family of chaperones that use an ATP-driven conformational cycle to recognize misfolded proteins, promote refolding, and prevent/resolubilize protein aggregation^1^. Hsp70s contain two highly conserved domains, the nucleotide-binding domain (Hsp70_NBD_) and the substrate-binding domain (Hsp70_SBD_) (Fig. E1a). The Hsp70 ATP/ADP cycle is regulated by two distinct protein families: Hsp40s, which induce ATP hydrolysis, and nucleotide exchange factors (NEFs), which facilitate the ADP/ATP exchange^25,26^. One of these NEFs is Bag1 and was initially identified as a binding partner of the anti-apoptotic protein Bcl-2 (Bcl-2 associated anthanogene 1)^27^. Bag1, which has been shown to interact with Hsp70^28,29^, is involved in a wide range of cellular pathways, including proteostasis, apoptosis, DNA transcription, proliferation, and neuronal homeostasis^27,29–32^. Bag1 contains UBL (Bag1_UBL_) and BAG (Bag1_BD_) domains (Fig. E1a); it interacts with the 26S proteasome through Bag1_UBL_ and with Hsp70_NBD_ through Bag1_BD_^16^.

Although the mechanism by which the proteasome degrades ubiquitinated proteins has been structurally elucidated^7,8^, it remains uncertain how the 26S proteasome, assisted by Bag1, translocates Hsp70-clients into the CP in a ubiquitin-independent manner. Here we report cryo-EM structures of the Bag1-bound 26S proteasome. These structures reveal that Bag1 plays a key role in the structural and functional regulation of the 26S proteasome. Our results demonstrate that Bag1 not only physically links Hsp70 to the proteasome through Rpn1, thus facilitating delivery of client proteins to the proteasome, but also triggers a series of conformational changes within the proteasome RP. These conformational changes serve to degrade client proteins without ubiquitination or ATP hydrolysis, thus revealing a previously undescribed mechanism of proteasomal degradation.

## Results

### The Hsp70-cochaperone Bag1 interacts with the proteasome subunit Rpn1

We first investigated, using size exclusion chromatography (SEC), whether Bag1 interacts directly with any of the three Ub receptors of the proteasome (Rpn1, Rpn10 and Rpn13 subunits). Complex formation was observed between Bag1 and Rpn1 through the Bag1_UBL_ (Fig. 1a and Fig. E1b), but not with Rpn10 (Fig. E1c) or Rpn13 (Fig. E1d). Rpn1 is composed of a N-terminal domain (Rpn1_NTD_), a C-terminal domain (Rpn1_CTD_) and a toroidal domain (Rpn1_TD_) (Fig. E1a). Rpn1_TD_ contains two Ub binding sites (T1 and T2) that interact with Ub chains and UBL domains, respectively (Fig. E1e)^33^. Deletion of the N-terminal Rpn1(1-260), termed Rpn1ΔNTD had no impact on Bag1 or Bag1_UBL_ binding, indicating an interaction of Bag1with Rpn1_TD_ (Fig. E1f). Furthermore, formation of a ternary complex consisting of Hsp70, Bag1 and Rpn1 was observed (Fig. 1b), despite of no direct interaction between Hsp70 and Rpn1 (Fig. E1g). Isothermal titration calorimetry (ITC) experiments showed that Bag1 had a higher affinity for Hsp70_NBD_ (K_D_ ≈ 50 nM) than for Rpn1 (K_D_ ≈ 500 nM) (Fig. E2a-c). Titration of Bag1 with an equimolar mixture of Hsp70 and Rpn1 resulted in the formation of the ternary complex (Fig. E2d), confirming that Bag1 is capable of concurrently binding to Hsp70 and Rpn1, as demonstrated by SEC. Hsp70 in the ternary complex retains the binding ability of unfolded proteins, as it is observed in SEC analysis in the presence of a Hsp70 client, reduced and carboxymethylated lactalbumin (RCMLA)^34^ (Fig. 1b). Thus, Bag1 recruits the client-bound Hsp70 to the proteasome subunit Rpn1.

**Fig. 1.**
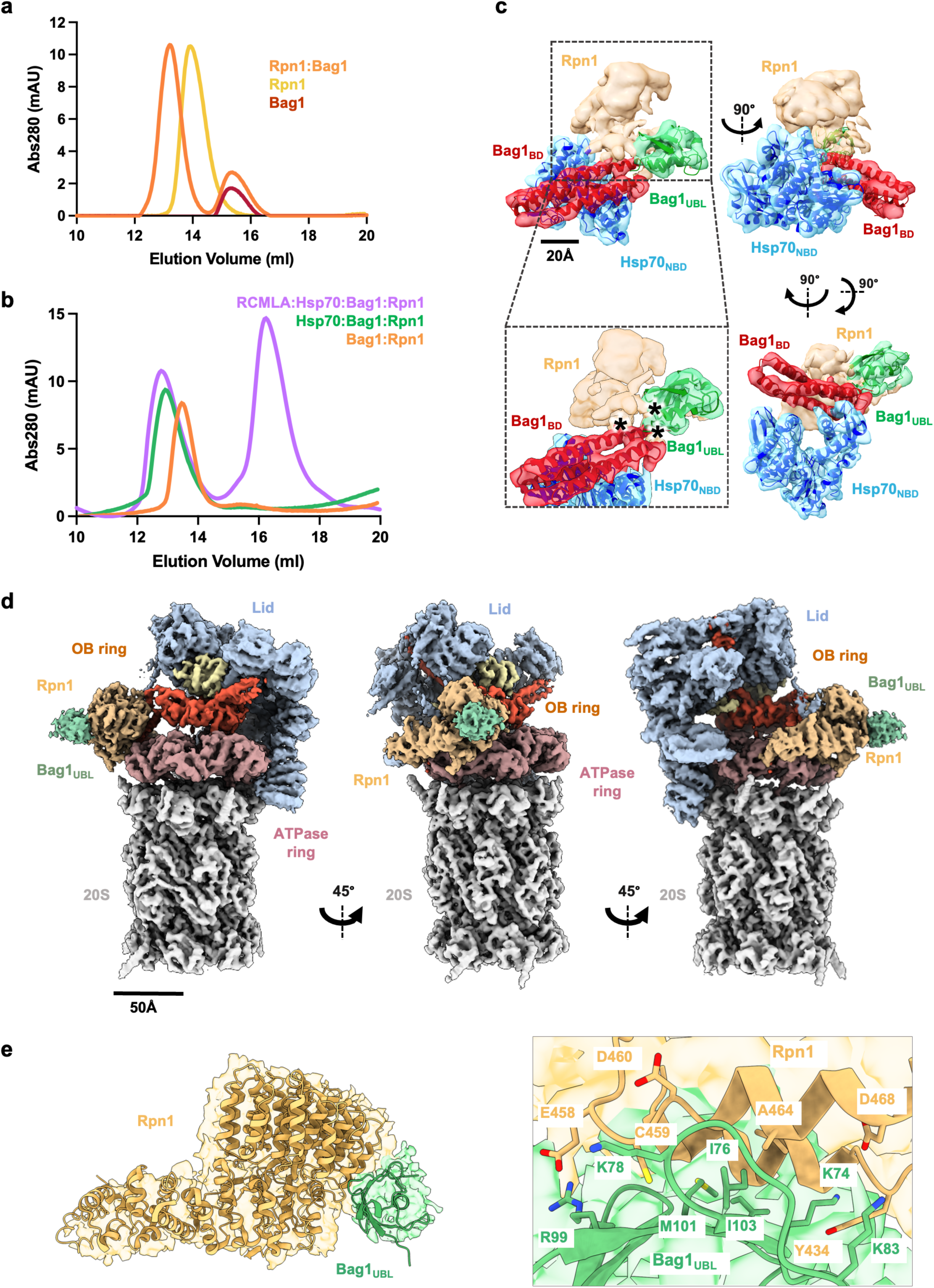
Bag1 interaction with the 26S proteasome and the chaperone Hsp70. (**a**) SEC analysis of Bag1 interaction with the proteasome subunit Rpn1. The shift in the elution profile of the sample containing both Bag1 and Rpn1 (orange) indicates the formation of a complex compared to Bag1 (red) and Rpn1 (goldenrod) alone. (**b**) SEC analysis of different combinations of Hsp70, Rpn1, Bag1 and a model substrate RCMLA. The sample containing Hsp70, Rpn1 and Bag1 (green) elutes prior to Bag1:Rpn1 complex (orange), showing a formation of a ternary complex. Upon addition of RCMLA to the ternary complex (purple), a shift in the elution peak was observed, showing that the model substrate interacts with the ternary complex of Hsp70:Bag1:Rpn1. (**c**) Different views of the cryo-EM map (4.8 Å resolution) of the Hsp70_NBD_:Bag1:Rpn1 ternary complex. AlphaFold prediction^37^ of Hsp70_NBD_ (blue) and full-length Bag1 (Bag1_BD_ in red and Bag1_UBL_ in green) are docked into the final map. The remaining density, which is presumably attributed to part of Rpn1, is colored in wheat. Bag1 interfaces to the putative Rpn1 density are indicated with black asterisks. **(d)** Cryo-EM reconstruction of the Bag1-bound 26S proteasome in S_BAG1_ (EMDB:52097) at 3.8 Å resolution. Only the UBL domain of Bag1 (Bag1_UBL_) is observed, with the BAG domain missing in the map. Colors are as follows: CP (white), ATPase domain of Rpts (rosy brown), OB domain of Rpts (orange), Rpn1 (beige), Bag1_UBL_ (light green), Rpn11 (light yellow), Lid (light blue). **(e)** Binding of Bag1_UBL_ to the T2 site of Rpn1 in the proteasome. The inset shows contacts between Rpn1 and Bag1_UBL._

### Structural characterization of the Hsp70_NBD_:Bag1:Rpn1 complex

Next, we analyzed by cryo-EM the structure of the ternary complex formed by Hsp70, Bag1 and Rpn1 (Hsp70:Bag1:Rpn1). To reduce the structural heterogeneity associated with the two Hsp70 domains^35^, we used only the Bag1 interacting domain of Hsp70, Hsp70_NBD_, thereby generating the Hsp70_NBD_:Bag1:Rpn1 complex, which was further stabilized with the crosslinker bis(sulfosuccinimidyl)suberate (BS3) (Fig. S1). After 3D classification, we obtained two classes of the ternary complex with different Bag1_UBL_ orientations (Fig. 1c, Figs. S2 and E3a-c). The interaction between Hsp70_NBD_ and Bag1_BD_ is identical to that observed in the crystal structure of Hsc70_NBD_:Bag1_BD_^36^ (Fig. E3d). The remaining density, likely corresponding to Rpn1, was too weak to be identified unambiguously due to the structural flexibility. To address this issue, we performed cross-linking mass spectrometry (XL-MS) analysis of the full-length Hsp70:Bag1:Rpn1 complex. Thereby we identified multiple inter-protein crosslinks between the three components, as well as intra-protein crosslinks within the individual proteins (Fig. E4a and Extended Table 1). Notably, both Hsp70_NBD_ and Hsp70_SBD_ were predominantly crosslinked to Rpn1_TD_, with fewer crosslinks observed with Rpn1_NTD_, leading us to conclude the remaining density in the cryo-EM map of the ternary complex corresponded to Rpn1. Both Bag1_BD_ and Bag1_UBL_ domains interact with the putative Rpn1 density, although Bag1_UBL_ engages with a larger contact surface (see black asterisks in Fig. 1c). There are no significant interactions observed between Hsp70_NBD_ and Rpn1, or Hsp70_NBD_ and Bag1_UBL_. Consistent with our structure, alphaFold-Multimer predictions^37^ of the ternary complex showed that Bag1 interacts with Rpn1_TD_ via Bag1_UBL_ (Fig E4b-e). In these predictions, the interaction between Rpn1 and Bag1_UBL_ is mediated by a hydrophobic patch around Bag1_UBL_ Ile76, which is reminiscent of the UBL binding of other proteasome binding proteins (Fig. E4d)^23,33,38,39^. The predicted model of Hsp70_NBD_:Bag1:Rpn1, based on our cryo-EM reconstruction, reveals that Rpn1_TD_ is sandwiched between Bag1_BD_ and Bag1_UBL_ (Fig. 1c and Fig. E4b-c).

### Cryo-EM structures of the Bag1-bound 26S proteasome

Having identified Rpn1 as the Bag1 binder, we next performed cryo-EM analysis of the Bag1-bound 26S proteasome stabilized with BS3 (Figs. S3 and S4). After the 3D classification, particles were separated into five conformational states (Fig. S3): three with an extra mass interacting with Rpn1 and two with no extra density bound to the proteasome. While the S_A_ state predominantly appeared in the control dataset of the human 26S proteasome^40^, the S_D_ state was more abundant in particles lacking the extra density in our dataset. In contrast, the three reconstructions with an additional density exhibit previously undescribed conformational states, which we termed S_BAG1_, S_BAG2_, and S_BAG3_ (Fig. S3 and S5). These three reconstructions had overall resolutions of 3.5∼3.6 Å (Figs. S3 and S4c). The observed extra mass in the S_BAG_ structures is associated with the T2 binding site^33,39^ of Rpn1_TD_, and was attributed to Bag1_UBL_ (Fig.1d-e and Fig. E1e), whereas the rest of the Bag1_BD_ density was not observed, probably due to its structural flexibility (see below). Subsequently, atomic models of the 26S:Bag1_UBL_ complex in the S_BAG_ states were generated (Fig. S5).

### Dynamic conformational alternation of the AAA+ ATPases induced by Bag1-binding

Comparison of the S_BAG_ states with previously reported EM maps showed significant dissimilarities in the interior of the RP but not in the CP (Figs. 2-4 and Figs. E5 and E6)^39,41^. In the S_BAG_ conformations, the positions of pore-2 loop of Rpt2, Rpt4, and Rpt5 are exceptionally closer to the CP α ring compared to the canonical structures, which will be discussed below. Of the conformations reported to date, the Rpt2 pore-2 loop is only located at the bottom of the staircase in the S_D4_ state (Fig. E7a,b), prompting us to use the S_D4_ structure as a reference for comparing the S_BAG_ structures. In the S_BAG1_ state, the lid complex tilts approximately ∼ 17.7° anticlockwise and shifts by ∼19.8 Å towards the Rpn1 N-terminus compared to the S_D4_ state (Fig. E6a). Rpn1 changes its position, shifting ∼15.0 Å and rotating ∼14.7 ° towards the interface connecting the ATPase and OB domains of both Rpt1 and Rpt2 (Fig. 2a,c,f). The Rpn1_TD_ lies in close proximity to the Rpt1 OB domain, and the Rpn1_CTD_ makes contact with the Rpt2 ATPase domain (Fig. 2c). Despite subtle conformational changes in Rpn1 (Fig. E6c,d), more drastic conformational changes are observed in the AAA+ ATPases (Figs. 2b,d-g, Fig. E6e, Fig. S6). The ATPase ring no longer retains its spiral staircase conformation but adopts an asymmetric hexameric shape with a significant distortion (Fig. 2g,3b, Fig. E7a and Supplementary Video 1). This deformation is caused by drastic conformational changes of all six Rpt subunits, most pointedly with Rpt4 shifting off-center (Fig. 3c,d,f). The fine architecture of the central channel, where the substrate-binding pore loops are arranged in a staircase, becomes disrupted and deformed, resulting in the formation of a substantial internal cavity at the center of the ring (Fig. 2b,3b,4a Fig. E8c and Supplementary Video 2). In the S_BAG1_ structure, the Rpt subunits are positioned at random heights rather than the ordered spiral arrangement seen in the canonical 26S structures (Fig. E7a). Similar distortions including a large cavity were observed in the other two conformations (S_BAG2_ and S_BAG3_) (Figs. E8, S6). Altogether, the S_BAG_ structures represent novel conformations, significantly departing from all proteasome states known to date.

**Fig. 2.**
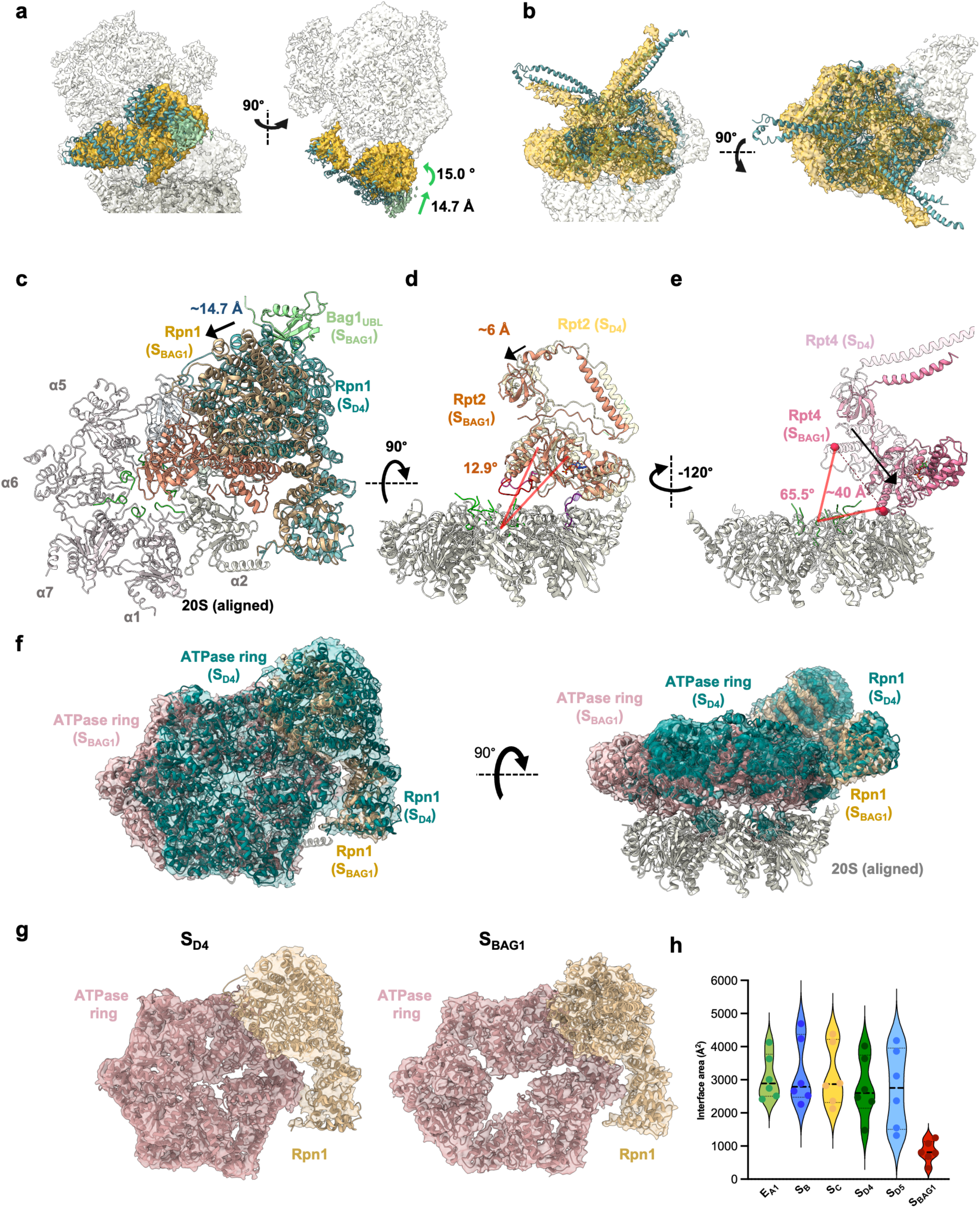
Dynamic conformational changes of the 26S proteasome induced by Bag1. **(a,b)** Structural comparison of the cryo-EM reconstruction of the 26S proteasome in S_BAG1_ (EMDB: 52097 in goldenrod) with the S_D4_ state (PDB: 7W3K in teal), focusing on Rpn1 **(a)** and ATPase ring **(b)**. Bag1_UBL_ is shown in light green and the rest of densities are shown in light grey. The Changes in shift (Å) and angle (°) are indicated. **(c-e)** Comparison of individual subunits in S_BAG1_ and S_D4_ (PDB: 7W3K) states. Structural differences in Rpn1 **(c)**, Rpt2 **(d),** and Rpt4 **(e)** are shown. Two structures are aligned to the CP α ring. The atomic model of the 20S CP is shown in white. Rpn1 is shown in beige for S_BAG1_ and in teal for S_D4_ **(c)**. Rpt2 and Rpt4 in the S_BAG1_ are depicted in dark salmon and pale violet red, respectively, while the structures in the S_D4_ are shown in transparent **(d,e)**. **(f)** Superimposition of the S_BAG1_ (EMDB: 52097, PDB: 9HEU) and S_D4_ (EMDB: 32283, PDB: 7W3K) Rpn1 and ATPase ring structures. The two cryo-EM structures are aligned to the CP α ring. In S_BAG1_, the ATPase ring (rosy brown) protrudes outward relative to the 20S CP, compare to the S_D4_ (blue green). Rpn1 (beige) shifts and rotates toward the ATPase ring. Atomic models for each map are shown. **(g)** Structural comparison of the ATPase ring (rosy brown) and Rpn1 (beige) in the S_BAG1_ (left) and S_D4_ (right) reveals that the ATPase ring in S_BAG1_ is deformed and creates a large cavity at the center. **(h)** Averages of the contact area between the AAA+ domains of adjacent Rpt subunits in different conformational states. Individual values for each structure are shown in dots and the median with a black dashed line. The S_BAG1_ has overall contact surfaces 3.5-fold smaller than the other conformational states.

**Fig. 3.**
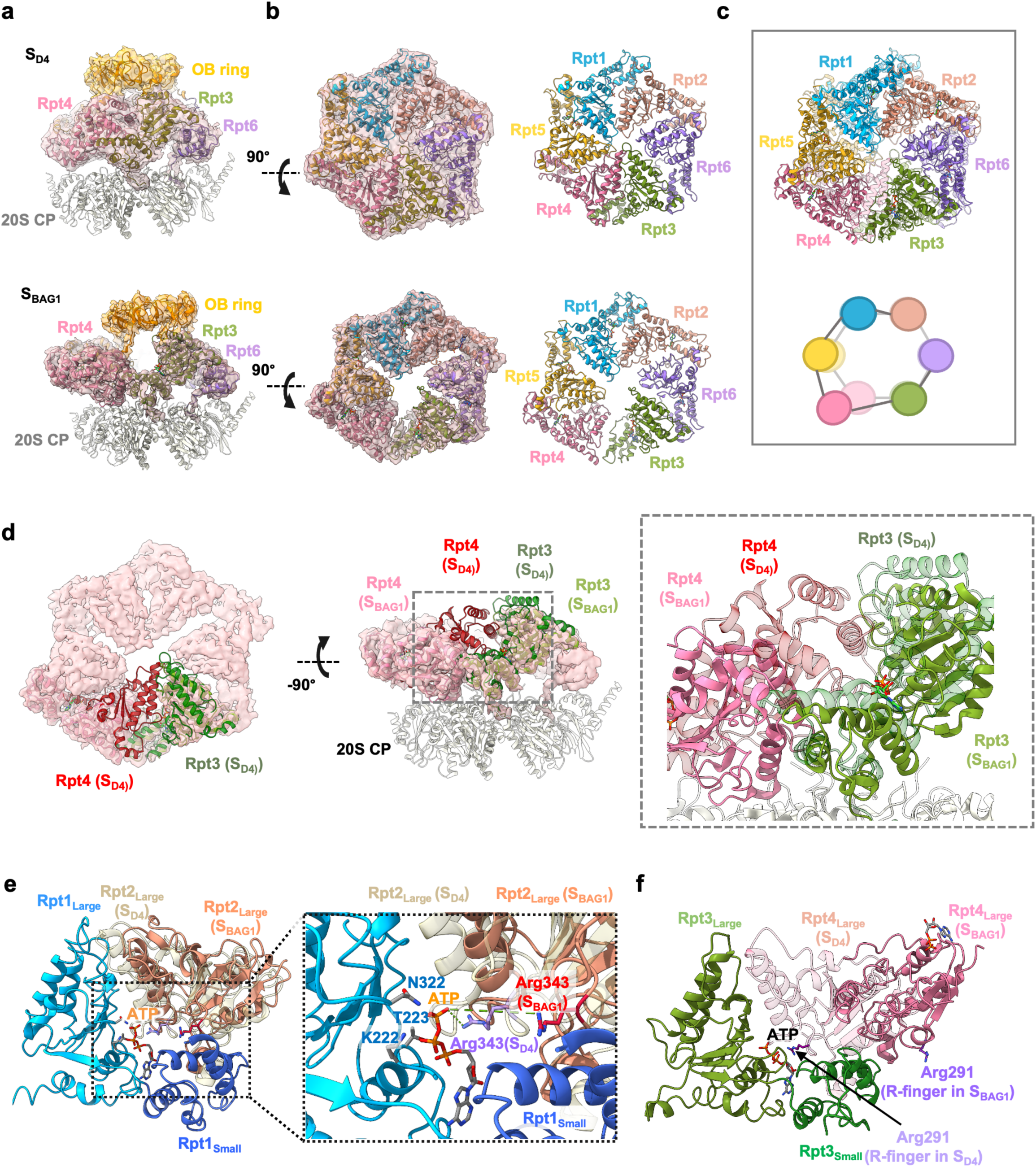
Deformation of the ATPase ring in S_BAG1_. **(a)** Cross-section of cryo-EM maps in S_D4_ state (upper panel) (EMDB: 32282; PDB: 7W3K) and Bag1-bound conformations in S_BAG1_ (S_BAG1_ PDB: 9HEU, EMDB: 52097), focusing on the interface between the OB and ATPase rings. Conformational changes in Rpt4 and Rpt3 (see schematic representation in **(c)**), result in a significant opening at the interface between the OB and ATPase domains. Cryo-EM densities are shown in transparent. **(b)** Cryo-EM segmentation of the ATPase ring in S_D4_ and S_BAG1_ (PDB: 9HEU; EMDB: 52097). A large cavity is observed in the middle of the ATPase ring in all three Bag1-bound conformations, while the ATPase ring is tightly packed in S_D4_. Atomic models of the subcomplexes, and individual Rpt subunits are colored as followings; OB ring (orange), 20S CP (white), Rpt1 (light blue), Rpt2 (salmon), Rpt6 (purple), Rpt3 (green), Rpt4 (hot pink), Rpt5 (yellow). **(c)** Schematic representation of the movement of the ATPase ring from a top-view. Structural models in S_BAG1_ and S_D4_ are aligned with the 20S CP and only the ATPase rings are shown with the same color code in Fig. 3b. The S_D4_ is shown in transparent. **(d)** Structural comparison between S_BAG1_ and S_D4_ focusing on the conformational change of Rpt3 (olive) and Rpt4 (hot pink). The S_D4_ structure is shown in transparent. **(e)** Structural comparison of the Rpt2 large domains at the interface of Rpt1 between S_BAG1_ (PDB: 9HEU) and S_D4_ (PDB: 7W3K). Two structures are superimposed with Rpt1 large domain (light blue). Only Rpt1 subunit in S_BAG1_ is shown (large and small domains in light blue and in blue, respectively). Rpt2 large domains in S_BAG1_ (salmon) and S_D4_ (light yellow) are shown for comparison. Rpt2 position is shifted and Arg(R)-finger Arg343 is positioned far from ATP in S_BAG1_, implying a lack of ATPase activity. Detailed view is shown in the dotted square. (**f**) Structural comparison of the Rpt4 large domain at the interface of Rpt3 between S_BAG1_ and S_D4_. Two structures are superimposed with Rpt3 large domain (olive). Only Rpt3 subunit in S_BAG1_ is shown (large and small domains in olive and green). Rpt4 large domains in S_BAG1_ (hot pink) and S_D4_ (light pink) are shown for comparison. Rpt4 large domain in S_BAG1_ shifts by ∼40 Å and rotates 65.5° in comparison to SD4 (Fig. 3d). Rpt4 Arg-finger Arg291 is highlighted.

**Fig. 4.**
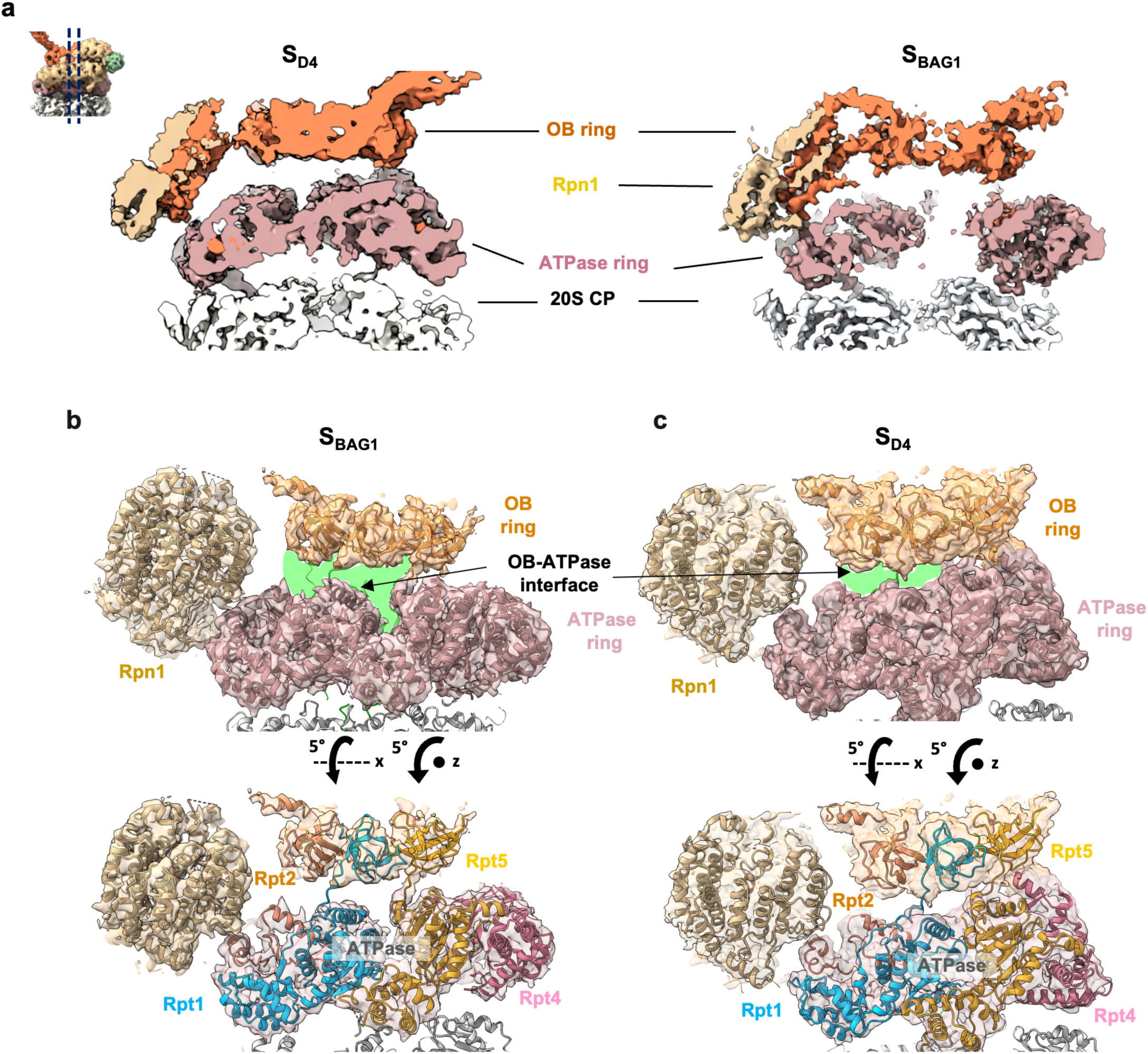
**(a)** Cross-section of cryo-EM map of the proteasome in S_BAG1_ and S_D4_ focusing on the interface between the ATPase and CP rings. Rpn1 (tan), OB ring (orange), ATPase ring (rosy brown), and CP (white) are colored separately, as indicated. In S_BAG1_, the central channel is deformed and a large cavity is observed on top of the CP gate, whereas the interior of the ATPase ring is packed in S_D4_. **(b)** In S_BAG1_, the atypical positioning of the ATPase subunits creates a large cleft (highlighted in light green) between the OB (orange) and ATPase (rosy brown) rings. The structure contrasts with the S_D4_ structure (EMDB: 32283) (PDB: 7W3K) in **(b)**. The atomic models of Rpt1, Rpt4 and Rpt5 are colored in sky blue, pink and goldenrod, respectively

As described above, the 3D reconstructions of the S_BAG_ states only reveal the presence of Bag1_UBL_ bound to Rpn1_TD._ Therefore, we investigated whether this interaction alone was enough to generate such large conformational changes in the proteasome. However, 3D reconstructions of the 26S in the presence of the isolated Bag1_UBL_ or Bag1_BD_ domains did not display such conformations and instead exhibited the S_A_ state (Fig. S7). This indicates that, although only Bag1_UBL_ is visible in the S_BAG_ structures, full-length Bag1 is needed to induce the conformational changes that lead to the loss of the central channel function in the 26S proteasome.

The highly conserved packed-interfaces between adjacent ATPase subunits, which are regarded as ‘rigid bodies’ in the known AAA+ ATPases (e.g. S_A_-_D_, S_D4,_ E_A-D_)^8,39^, are widely open in S_BAG1_, with contact surface areas ∼1000 Å^2^, dramatically smaller than those observed in any of canonical interfaces with average contact area of 2996 Å^2^ (Figs. 2h, 3e,f, and Fig. E7c). Across all interfaces, the Arg finger motifs, crucial for the interaction with the ATP γ-phosphate of the neighboring subunit, are not directed towards the nucleotides in the Bag1-bound 26S proteasome structures (Figs. 3e,f, Fig. E7e). In particular, Rpt4, rotated by ∼65.5° and shifted by ∼40 Å, interacts neither with the Rpt3 nucleotide nor with its large domain, and maintains only limited contact with the Rpt3 small domain, which covers 331.5 Å^2^ (Fig. 3f). The S_BAG1_ map shows nucleotide density in all six nucleotide-binding pockets; ATP density for Rpt1 and Rpt3, and ADP density for Rpt2, Rpt6, Rpt4 and Rpt5 pockets (Fig. E7f). However, the disengagement of the Arg fingers and the large distance at the interfaces would indicate lack of ATPase activity for the Bag1-bound 26S proteasome. Indeed, we demonstrated that ATPase activity decreased upon full-length Bag1 titration (Fig. 5a). Although a previous study showed that UBL binding does not stimulate the ATPase activity^42^, Bag1_UBL_ also modestly reduced ATPase activity, while Bag1_BAG_ had no effect. Given that Bag1_UBL_ does not induce the ATPase deformation, the unique N-terminal segment of Bag1, which is rich in charged residues, likely contribute to the modulation of ATPase activity. Thus, Bag1 binding deforms the interfaces of the Rpt subunits and blocks the ATPase activity of the 26S proteasome.

**Fig. 5.**
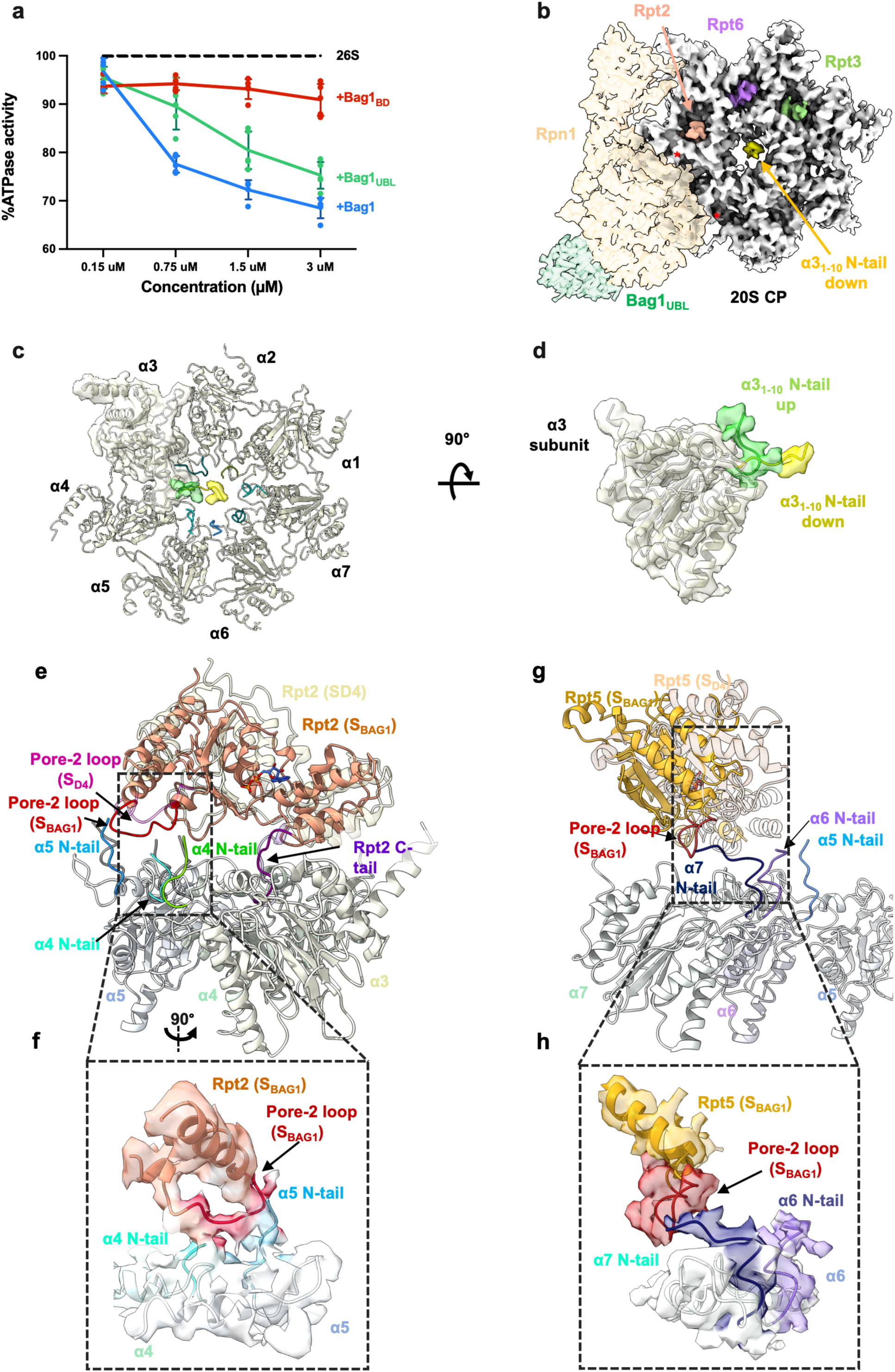
CP gate opening mechanism in the Bag1-bound 26S proteasome. **(a)** ATPase activity of the 26S proteasome (black dash line) upon Bag1 (blue) Bag1_BD_ (red) or Bag1_UBL_ (green) titration. Whereas Bag1_BD_ shows no effect, full-length Bag1 and Bag1_UBL_ decrease 26S ATPase activity. The data represent the mean ± SD for n=5 independent experiments (represented with dots). **(b)** Insertion of Rpt C-terminal tails into CP a ring pockets. The RP–CP interface and insertion of Rpt C-terminal tail into the α-pockets of the CP in S_BAG1_ are shown. The cryo-EM density of the CP is shown in white, whereas the C-terminal tails of Rpt2, Rpt3 and Rpt6 are colored in dark salmon, green and purple respectively. Empty pockets are indicated with red asterisks. The EM density of the N-terminal tail of α3 in the ‘down’ state is shown in yellow. **(b,c)** N-terminal tail of α3 exhibits ‘up’ and ‘down’ states. The ‘up’ conformation (light green) corresponds to the ‘open’ gate, while the ‘down’ conformation (yellow) represents the ‘closed’ gate. Side view of the α3 subunit highlights the movement of the N-terminal tail **(c)**. **(d)** The pore-2 loop of Rpt2 moves lower towards the CP gate, by approximately 2 Å distance and interacts with N-terminal tails of α4 and α5. **(e)** Zoom-in of the cryo-EM density of the Rpt2 pore-2 loop and the N-terminal tails of α4 and α5. **(f)** The pore-2 loop of Rpt5 moves lower towards the CP gate, approximately 4 Å distance and interacts with N-terminal tails of α6 and α7. **(g)** Zoom of the cryo-EM density of the Rpt5 pore-2 loop and the N-terminal tails of α6 and α7. For **(d-g)** comparison with S_D4_ (PDB: 7W3K) was used.

### Mechanism of gate opening by Bag1

Surprisingly, all S_BAG_ structures show open gate conformations with apertures of varying diameter, similar to previous observations in the substrate-bound 26S structures^7,8^ (Fig. 5b and Fig. E8f). In S_BAG1_ and S_BAG3_, the N-terminal tails of all α subunits, which form the 20S CP gate, adopt an open conformation except for the α3 N-terminus (Figs. 5c,d and Fig. E8f), which shows two conformations; perpendicular (up) or parallel (down) to the CP plane (Fig. 3d), whereas the gate in S_BAG2_ is completely open (Fig. E8f). The coordinated insertion of the five Rpt C-termini (Rpt2, Rpt3, Rpt5, Rpt1 and Rpt6) into the α pockets are critical for gate opening, as observed in S_D4_^7,40,43^ (Fig. E8f). In the three S_BAG_ structures, only densities of the C-termini of three subunits (Rpt2, Rpt3, and Rpt6) are detected in the α pockets, placing the N-termini of α1, α2, and α3 in the open-gate conformation (Fig. 5b and Fig. E8f and Supplementary Video 3). In contrast, the Rpt1 and Rpt5 C-termini, which regulate the α4 and α5 N-termini, are not observed. Instead, the N-termini of the α4 and α5 subunits are held up by their interaction with the Rpt2 pore-2 loop (Fig. 5e-f and Supplementary Video 4). Additionally, the pore-2 loop of Rpt5 supports the α7 N-terminus, which further interacts with the α6 N-terminus (Fig. 5g-h and Supplementary Video 4). The conserved pore-2 loops (Fig. E9a) in the previous substrate-bound proteasome structure form a spiral staircase below the pore-1 loops, with both loops intercalating into the processing polypeptide within the central channel^7,8,39^. The pore-2 loops in the Bag1-bound 26S structures, especially Rpt2, Rpt4 and Rpt5, position much closer to the CP compared to the canonical AAA+ structures (Fig. 5e-h, Figs. E7a-b and Supplementary Video 4). Although the Rpt4 pore-2 loop also lowers its position, its density disappears above the α pocket between α7 and α1, thus not allowing us to decipher its role (Fig. E9b). Finally, the shift of the Rpt2 pore-2 loop towards the CP gate is coupled with a global structural change of the Rpt2 subunit, which is induced by a conformational alternation of Bag1-bound Rpn1. Deformation of the ATPase rings impairs the pore loop function for substrate translocation, but instead provides a novel function, enabling substrate entry into the CP by gate opening (Fig. 5e-h and Supplementary Video 4).

### Mechanism of Bag1-induced proteasomal degradation

Bag1-induced deformation of the ATPase ring lowers the individual ATPase subunits, placing them closer to the CP ring. Besides, the respective position of the OB ring and DUB Rpn11 exhibits a slight change (Figs. E6b and E10a,b). These conformational changes create a gap of approximately 20 Å between the OB ring and the ATPase ring (Fig. 4a-b, Fig. E10c-h and Supplementary Video 5). All OB-ATPase interfaces are loosened compared to those of the canonical proteasome structures, particularly for Rpt4 and Rpt5, which are off-centered and largely protrude from the CP cylinder (Fig. E10c-d). Notably, the wide OB-ATPase opening is interconnected to the prominent cavity at the center of the ATPase rings, reaching the CP gate (Fig. 4). The architecture of this cavity suggests a role as a substrate chute for direct protein degradation. To understand the biological role of Hsp70:Bag1 binding to the 26S proteasome, we fitted the structural model of the Hsp70_NBD_:Bag1:Rpn1 complex (Fig. 1c) into the 26S:Bag1 complex, as well as the AlphaFold prediction of ADP-bound Hsp70 full length structures. Notably, Hsp70_SBD_ can be placed without clashing with the ATPase rings of the proteasome, and the Hsp70 hydrophobic patch that mediates substrate binding points towards the OB-ATPase gap created by Rpt4 and Rpt5 (Fig. 6a). This structural model suggests an optimal transfer of the unfolded protein directly from Hsp70 to the 20S CP chamber for degradation.

**Fig. 6.**
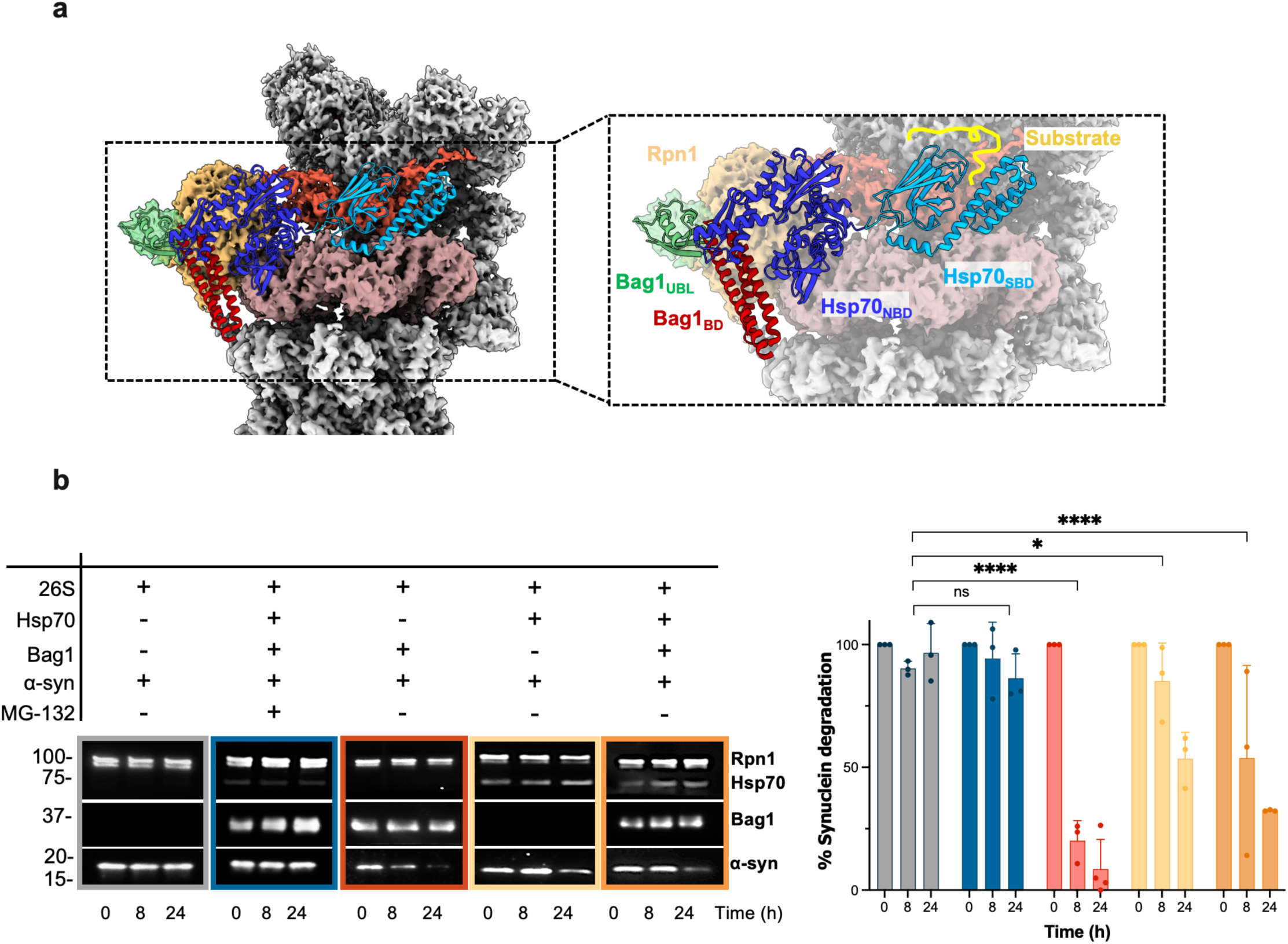
Structural basis for the unfolded protein transfer. **(a)** Structural model of the Hsp70-Bag1-bound 26S proteasome created based on Hsp70_NBD_:Bag1:Rpn1 complex (Fig. 1c) and the 26S:Bag1 complex (Fig 1d) together with an AlphaFold prediction of the ADP-bound Hsp70 and Bag1 complex. Hsp70_SBD_ (dark blue) is positioned adjacent to the OB-ATPase cleft, indicating a direct transfer of unfolded proteins to the 20S CP for degradation. **(b)** Summary of western blot results (left panel) analyzing proteasomal degradation of α-synuclein in the absence of ATP at 0, 8, and 24 hours. Statistical analysis (right panel) reveals that Bag1 alone (red) and with Hsp70 (orange) significantly enhance synuclein degradation compared to the proteasome alone (grey), while Hsp70 alone (yellow) shows stronger effects at later times. MG-132, as expected, inhibits degradation (dark blue). Data (n=4-5) analyzed via two-way ANOVA (*p=0.0402, ****p<0.0001).

The disrupted interface of the nucleotide binding pockets accounts for the reduced ATPase activity of the proteasome upon Bag1 binding (Fig. 5a). To address whether Bag1 influences the degradation of unstructured proteins, we used α-synuclein (α-syn) as a model substrate. α-syn, an IDP, is highly expressed in neurons, particularly in presynaptic terminals^44^. An *in vitro* substrate degradation assay using α-syn showed an enhancement of degradation in the presence of Bag1 (Fig. 6b and Fig. S9). Interestingly, when combined with Hsp70, Bag1 moderated the degradation rate of α-syn (Fig. 6b and Fig. S8). Positioned in the proximity of the opening between OB and ATPase rings, Hsp70 slowed down α-syn degradation, suggesting that it selectively restricts access of unstructured substrates to the proteasome. Thus, Bag1 can enhance degradation of unstructured proteins in an ATP-independent manner by promoting conformational change within the proteasome in an ubiquitin- and ATP-independent manner.

## Discussion

In this work, we elucidate the structures of the Bag1-bound 26S proteasome, which exhibit striking differences from previously reported proteasome structures^7,8,39–41,43^. Particularly, the ATPase ring structures vary markedly from any known AAA+ ATPase structures^9^. Similar to observations of the proteasome structure in complex with DUB Usp14/Ubp6-bound and other UBL-containing cofactors^23,33,38,39^, Bag1 docks at the T2 site of Rpn1 through its UBL domain (Fig. 1d-e). Thus, Bag1 can potentially exclude the DUB Usp14, which plays a crucial role in trimming Ub chains, as it is not required for ubiquitin-independent degradation. The interaction between Bag1_UBL_ and Rpn1 appears to reorientate Rpn1 towards Rpt2, promoting significant structural deformation within the ATPase ring. Displacement of individual Rpt subunits, particularly Rpt4, which is significantly rotated and shifted off-center, results in the formation of a wide cavity at the center the ATPase ring (Fig. 2,3), located above the now open CP gate (Fig. 4a and Supplementary Video 2). The ATPase ring alters its conformation from a canonical, spiral staircase conformation to an asymmetric hexameric ring (Supplementary Videos 1 and 5). The conserved architecture of the AAA+ hexamers and tightly closed subunit interface are the basis for converting the energy of ATP hydrolysis into mechanical force^9^. However, in the Bag1-bound 26S structures, the disrupted interfaces of the nucleotide binding pockets lead to a loss of ATP hydrolytic activity (Fig. 5a).

During ATP-dependent ubiquitinated substrate degradation, the pore loops lining the central channel of the AAA+ ATPases normally make direct contact with the substrate polypeptides. The pore-2 loops form a spiral staircase beneath the pore-1 loops, with both loops intercalating into the processing polypeptide within the central channel^7,8,39^. The conserved aromatic residue of the pore-1 loop enhances the grip on the substrate, while the pore-2 loop supports this function using its aromatic residue surrounded by charged residues^9^. In contrast, in the Bag1-bound structures, the pore-2 loops of Rpt2 and Rpt5 are involved in 20S gate-opening together with the C-terminal tails of Rpt2, Rpt3 and Rpt6 (Fig. 5 and Supplementary Videos 3,4), providing insights into a novel role of the pore-2 loops. These arrangements are reminiscence of the mitochondrial AAA+ protease AFG3L2, as its pore-2 loop at the bottom of the staircase is positioned lower than in canonical structures, and contacts the center of the proteolytic chamber^45^.

Cooperation with a certain set of cochaperones enables linking of the Hsp70 chaperones to the 26S proteasome for degradation of unfolded proteins^2^. Bag1, which interacts with Hsp70_NBD_, is able to recruit Hsp70-bound clients to the 26S proteasome through its UBL domain^16^. By combining our cryo-EM structures of the Hsp70_NBD_:Bag1:Rpn1complex and the Bag1-bound 26S proteasome, we created a structural model of the Hsp70-Bag1-26S proteasome complex (Fig. 6a). This model suggests that the Hsp70_SBD_, to which unfolded proteins are associated, could be positioned near the expanded interface between the OB and ATPase rings, which connects to a large ATPase cavity and the opened CP gate. Thus, our structures illustrate how Bag1 could assist the 26S proteasome in enhancing substrate transfer from Hsp70, thereby facilitating substrate entry into the 20S CP. The deformed ATPase ring presumably allows the 26S proteasome to degrade unfolded proteins directly in a ubiquitin- and ATP-independent manner. Interestingly, our degradation assay revealed faster degradation in the presence of Bag1 alone compared to when both Hsp70 and Bag1 are present. The structural model suggests that Hsp70_SBD_ can occlude the OB-ATPase opening, potentially helping to prevent non-specific protein degradation. Additionally, considering that Bag1 has a higher affinity for Hsp70 than for Rpn1, a mechanism probably exists whereby client-bound Hsp70/Bag1 preferentially binds to the 26S proteasome. Taken together, our structural study illustrates a molecular mechanism by which unfolded proteins can be degraded by the 26S proteasome without the need of energy consumption (Supplementary Video 6) by the proteasome itself.

Cochaperone CHIP E3 ligase interacts with Hsp70_SBD_ and ubiquitinates the Hsp70-bound client proteins^46–48^. The ubiquitinated client is then recruited to the proteasome through its polyubiquitin chain and gets degraded in an ATP- and ubiquitin-dependent manner^48^. Although a direct interaction between CHIP and Bag1 has been reported^49^, it remains unclear whether Bag1 and CHIP operate simultaneously for unfolded protein degradation, as Bag1 reduces the ubiquitination efficiency of CHIP by altering the duration of Hsp70-client binding^50,51^.

Recent studies have shown that a subset of eukaryotic proteins is subjected to ubiquitin-independent proteasomal degradation^17–21,52^. Proteomics analysis and global protein stability assay identified nearly 2,000 proteins that are degraded by ubiquitin-independent proteasomal degradation, with some of these proteins specifically targeted only under stress conditions^18,22^. Many adaptor proteins such as midnolin, FAT10, or antizyme have been shown to assist direct protein degradation in an ATP dependent manner^22,23^. In our *in vitro* experiment using the mostly unstructured protein α-syn, we demonstrated that Bag1 can enhance α-syn degradation by 26S proteasome, suggesting that Bag1 may assist in direct recruitment of Hsp70 clients to the 26S proteasome independently of ubiquitination and ATP hydrolysis. This is a unique pathway, different from other ubiquitin-independent pathways assisted by adaptor proteins, and may be particularly important under ATP-deficient conditions, such as starvation. In addition, Bag1 has been reported to bind directly to multiple degradation targets^31,53–55^, raising the possibility of direct degradation by the 26S proteasome.

Bag1 plays a critical role in maintaining cellular homeostasis under stress conditions, where a rapid cellular response is likely required to prevent protein aggregation^56–58^. It has been reported that Bag1 protects neuronal cells from the toxicity of various amyloid proteins including α-synuclein^59–62^. Besides, Hsp70 interacts with IDPs even in the absence of ATP, preventing the amyloid formation^63–65^. It is tempting to speculate that in stress conditions, Bag1 could be upregulated similarly to what occurs with Hsp70 to facilitate the removal of unfolded proteins including amyloid-forming proteins, without the need of ubiquitin tagging and energy consumption. Taken together, this study describes a novel mechanism of proteasomal degradation, which not only advances our understanding of the mechanical function of the 26S proteasome, but also opens new avenues for therapeutic intervention in diseases associated with protein misfolding and aggregation.

## Methods

### Cloning, expression and purification of proteins

Human Rpn1, Rpn10 and Rpn13 genes were amplified from the cDNA library Human MTC Panel I (Clontech) and cloned into pPROEX-HTa vectors using In-Fusion® technique (Takara). Proteins were expressed in *E. coli* Rosetta (Rpn1) or C41 strain (Rpn10 and Rpn13). Expression was induced with 1 mM IPTG at the exponential phase (Rpn10) or using AutoInducible Medium (Rpn1, Rpn13) (CondaLab). To purify the proteins, cellular lysates were loaded onto HisTrap columns (Cytiva). After TEV protease cleavage, to remove the His_6_ tag, proteins were further purified using a Q column (Cytiva) and size exclusion chromatography (Superdex 200/75) and the resulting samples were stored at −80 °C in 20 mM Hepes pH 7.4, 150 mM KCl, 10 % glycerol.

Bag1 and Hsp70 were purified as previously described ^66^. Hsp70_NBD_, Rpn1ΔNTD, Bag1_UBL_ and Bag1_BD_ were purified as their full-length versions. Reduced Carboxymethylated Lactalbumin (RCMLA) were kindly provided by Fernando Moro (Universidad del País Vasco, Spain). Either HEK293T cells (purchased from the Helmholtz Center for Infection Research, Brunswick, Germany) or Expi293F cells (from the Netherlands Cancer Institute, Amsterdam, The Netherlands) were used for proteasome purification as described in ^67^. Approximately 18 g of cells were thawed and resuspended in 18 mL of 2X proteasome lysis buffer (100 mM Hepes pH 7.6, 10 mM DTT, 20 mM MgCl_2_, 10 mM ATP, 20% [v/v] glycerol), subsequently lysed and centrifuged at 20,000 rpm for 20 min at 4 °C in an SS-34 rotor. The sample was incubated with 3 µM of UBL-RAD23-GST and 1 mL of Glutathione Sepharose 4B (Cytiva) for 3 h, at 4 °C. After washing with proteasome lysis buffer, proteasomes were eluted with the same buffer supplemented with 5 mg/mL of UIM-HIS_5_ protein. The eluted sample was applied to a 15-30% (w/v) sucrose gradient (50 mM Hepes pH 7.6, 50 mM KCl, 5 mM DTT, 10 mM MgCl_2_ and 7.5 mM ATP and then sucrose to 15 or 30% [w/v]). The gradients were centrifuged at 33,000 rpm for 17 h at 4 °C in an SW40 rotor. Fractions showing peptidase activity were collected and flash frozen in liquid nitrogen and stored at −80 °C.

### Binding assays and SEC analysis

Protein mixtures were incubated for 20 min at 37 °C in a buffer containing 20 mM Hepes pH 7.4, 150 mM KCl, and 1 mM DTT. The samples were analyzed by SEC using either a Superdex 75 Increase 10/300 GL (Cytiva) or a Superdex 200 10/300 GL (Cytiva), depending on the complex size. The eluted fractions were further analyzed by SDS-PAGE. For the experiment with the quaternary complex, Hsp70 and RCMLA were first incubated for 1 h at 37 °C. Bag1 and Rpn1 were then added and incubated for another 20 min at room temperature. The sample was injected into the SEC column.

### Isothermal titration calorimetry (ITC) experiments

Proteins were exhaustively dialyzed against 20 mM phosphate buffer pH 7.4, 150 mM KCl. The final protein concentration was measured by UV-spectroscopy using the theoretical absorption coefficients at 280 nm (Abs_280_). ITC experiments were performed at 25 °C using a MicroCal PEAQ-ITC microcalorimeter (Malvern). Titrations were performed by stepwise injections of a 100-150 μM Bag1 solution into the reaction cell loaded with the other protein/s (16 μM Hsp70_NBD_, Rpn1 or both). Dilution heats were determined separately and subtracted from the total heat produced following each injection. Titration data were analyzed using the AFFINImeter-ITC software (http://software4science.com). For binary complexes, the data were fitted to a one-site binding model. For the ternary complex, independent binding of Rpn1 and Hsp70_NBD_ was assumed, and data were fitted simultaneously with binary complexes using the same binding parameters for each ligand across both complexes.

### Crosslinking and mass spectrometry (XL-MS) analysis

The Hsp70_NBD_:Bag1:Rpn1 complex was cross-linked with 3 mM bis(sulfosuccinimidyl)suberate (BS3, Thermo Scientific) in 20 mM HEPES pH 7.4, 150 mM KCl, 1 mM DTT (1 h, 4 °C), followed by quenching with 50 mM Tris pH 7.4. BS3-crosslinked samples were incubated in Laemmli sample buffer for 5 min at 95 °C and subjected to SDS-PAGE. The gel was stained with Quick Coomassie (Generon) and the bands corresponding to the complex were excised and subjected to automated reduction with TCEP, alkylation with chloroacetamide, and trypsin digestion using an OT2 robot (Opentrons), as described in Shevchenko et al ^68^. The resulting peptide mixture was speed-vac dried and re-dissolved in 0.1% formic acid for LC-MS/MS analysis by Ultimate 3000 nanoHPLC (Dionex) coupled to an Orbitrap Eclipse (Thermo) or to a Orbitrap Fusion Lumos (Thermo). The HPLC was equipped with a PepMap Neo C18 trapping column (300 µm x 5 mm; Thermo) and a PepMap RSLC c18 column (75 µm x 50 cm; Thermo). Solvent A and Solvent B were 0.1% formic acid in water and 0.1% formic acid in acetonitrile, respectively. Separation was performed at 50 °C at a flowrate of 250 nl/min under the following gradient: 4% B for 2 min, linear increase to 35% B in 68 min, linear increase to 50% B in 6 min, linear increase to 90% B in 4 min and 90% B for 10min.

The mass spectrometer was operated in DDA mode. Each acquisition cycle had a maximum duration of 3 sec and consisted of a survey scan (375-1250 m/z) at 120,000 resolution (FWHM) and up to 20 MS/MS scans at 30000 resolution (FWHM). Peptides with charges 2 to 6 were selected for fragmentation applying a dynamic exclusion window of 40 s. For peptide identification, raw MS data were converted to mgf files with Proteome Discoverer v2.5 (Thermo) that were used for a database search with MeroX 2.0 MeroX 2.0^69^ against a custom-made database containing the sequences of each protein. Search parameters were set as follows: trypsin as enzyme allowing 2 (K) and 2 (R) missed cleavages, BS3 as crosslinker, MS tolerance of 10 ppm and MS/MS tolerance of 20 ppm, carbamidomethylation of cysteines as fixed modification and oxidation of methionines as variable modification. Peptide identifications were filtered at an FDR of < 5% and a minium MeroX score of 30. Alternatively, the database search was conducted with XlinkX (Thermo) with the same database and search parameters except for the number of missed cleavages that was set to 3 and the FDR threshold was < 1%. Although conventional constraint distances allowed by BS3 are up to 25 Å^70^, a broader range (up to 35 Å) was applied to account for protein dynamics and conformational flexibility. The distances that fall within the BS3 constraints established in this work were displayed and represented using Chimera package. XL-MS data have been deposited to the ProteomeXchange Consortium via the PRIDE partner repository with the dataset identifier PXD058407.

### Proteasome activity

The chymotrypsin 20S activity was assayed using the model substrate succinyl-Leu-Leu-Val-Tyr-7-amino-4-methylcoumarin (Suc-LLVY-AMC) (Enzo). A 5 nM concentration of proteasome was preincubated with Bag1 (concentration range from 0.001 μM to 30 μM) for 20 min at 37 °C in peptidase buffer (50 mM Tris pH 7.4, 100 mM KCl, 1 mM ATP, 0.5 mM MgCl_2_, 1 mM DTT and 0.025 mg/mL BSA), and later incubated with the substrate (10-50 µM) for 1 h at 37 °C. The fluorescence was recorded using 350 nm excitation and 450 nm emission wavelengths every 30 s for 1 h in a SpectraMax ID3 Microplate reader (Molecular Devices). Proteasomal activity was measured as the slope of the curve, and was normalised between experiments considering the control without Bag1 as 0% and the highest activity as 100%. The normalised activity was fitted to a sigmoidal curve of the form A = 100·C/(C+K_D_), where A is the activity; C is Bag1 concentration and K_D_ is the apparent binding constant.

### ATPase activity assays

These assays were conducted following the standard protocol of BIOMOL GREEN Kit (Enzo). A constant final concentration of 50 nM proteasome was used for the samples when titrating Bag1 with molar ratios of 1:1, 1:2.5, 1:5 and 1:10. Proteins were mixed in ATPase buffer (20 mM HEPES pH 7.4, 40 mM NaCl, 5 mM MgCl2) for a final volume of 25 μl and incubated 10 min with 25 μl of substrate buffer (20 mM Tris pH 7.5, 5 mM MgCl2, 1 mM ATP). Then, the reaction was stopped with 100 μl of BIOMOL GREEN reagent, incubated 20-30 min at 37°C to allow development of the green color and the absorbance at 620 nm was measured.

### Substrate degradation assays

Mixtures with 350 nM of purified proteasome were incubated in the presence of different concentrations of Bag1 (1.4 μM), Hsp70 (1.4 μM), Hsp40 (1.4 μM), α-synuclein (0.35 μM) and MG-132 (50 μM). Samples were then incubated at 4 °C for 24 h, taking 5 μl at 0, 8 and 24 h. Then, samples were boiled 5 min at 95 °C, mixed with SDS-PAGE loading buffer, and loaded in 4-15% Mini-Protean precast gels (BioRad) for SDS-PAGE. Protein bands from SDS-PAGE were transferred to a PVDF membrane (Immobilon-P transfer membrane from Millipore) soaked in transfer buffer (25 mM Tris, 200 mM Glycine, Methanol 20 % v/v), 0.1 % (w/v) SDS) using a semi-dry transfer system (Trans-Blot Turbo from BioRad) at 25 V 1.0 A for 30 min. Membranes were then blocked with 3 % (w/v) non-fat dried milk in PBS-0.05 % Tween (PBST) for 1 h room temperature with gentle shaking. Then three washing steps of 10 min in PBST were performed to remove the blocking solution, and the membranes were later incubated with primary antibodies (all diluted in PBST) against Rpn1 (PSMD2 A11, Santa Cruz Biotechnology, 1:300 dilution), Hsp70_NBD_ (501043, PALEX,1:1000 dilution), Bag1 (α-Histidine tag coupled to horseradish peroxidase, Santa Cruz Biotechnology, 1:4000 dilution) and α-synuclein coupled to horseradish peroxidase (Santa Cruz Biotechnology, 1:100 dilution), 1 h RT with gentle shaking. Three washing steps of 10 min in PBST were performed before incubating with a secondary antibody against mouse IgG coupled to horseradish peroxidase (NA931, Cytiva, 1:10000 dilution) 1 h at RT with gentle shaking. The membranes were washed again 3 times and developed using Clarity Western ECL substrate (BioRad). The signal of the chemiluminescent bands was analyzed with Fiji.

### CryoEM data acquisition

Aliquots of the 26S:Bag1 complex were vitrified in a Vitrobot Mark IV (Thermofisher Scientific) at 4 °C and 100% humidity. Quantifoil Cu/Rh R2/2 300 mesh grids pretreated with poly L-lysine (Sigma Aldrich) were used. Grids without glow discharge were incubated for 90 s with 0.1 % poly L-lysine (w/v), then washed with water and dried ^71,72^. 14,079 images were acquired in a Titan Krios (EMBL, Grenoble, France) equipped with a K3 detector (Gatan Inc). A total dose of 37 e^-^/Å^2^ was applied to the images with nominal defocus ranged from −1 to −3 μm. The magnification was 81,000x, corresponding to a pixel size of 1.06 Å.

The Rpn1:Bag1:Hsp70_NBD_ complex was vitrified in a Vitrobot Mark IV (Thermofisher Scientific) at 4 °C and 100% humidity. Glow-discharged UltrAuFoil R1.2/1.3 300 mesh grids were used. 13,724 images were acquired in a Titan Krios (EMBL, Grenoble, France) equipped with a K3 detector (Gatan Inc). A total dose of 42 e^-^/Å^2^ was applied to the images with nominal defocus range from −1.4 to −3 μm and a magnification of 81,000x, corresponding to a pixel size of 1.06 Å.

The 26S proteasome samples in the presence of the isolated Bag1_UBL_ or Bag1_BD_ domains were vitrified in a Vitrobot Mark IV (Thermofisher Scientific). 3 μl of the samples were applied to freshly glow discharged Quantifoil Cu R2/2 200 mesh grids, washed with water and blotted. 6,685 and 5,051 images were acquired for 26S: Bag1_BD_ and 26S:Bag1_UBL_ samples, respectively, in a Titan Krios (Thermofisher Scientific) equipped with a Selectris energy filter operating at a slit width of 10 eV and Falcon4i detector (Thermofisher Scientific). The magnification was 105,000x, corresponding to a pixel size of 1.18 Å. A total dose of 40 e^-^/Å^2^ was applied to the images with nominal defocus range from −1 to −2 μm. Details of the data acquisitions are provided in Extended Table 2.

### Image processing and 3D reconstruction

Initial pre-processing of the 26S:Bag1 complex images and cleaning up of particles were performed using the Scipion 3.0 software platform (de la Rosa-Trevín et al., 2016). It began with movie frame alignment using MotionCorr2 ^73^, followed by CTF estimation by CTFFIND4 ^74^. 1,891,841 particles were picked by crYOLO ^75^. A total of 605,539 particles were identified after rounds of CryoSPARC 2D classifications and hereafter image processing was performed on standalone Relion-4.0 ^76^ or CryoSPARC ^77^. A further Relion 4.0 2D classification reduced the number of good particles to 542,965, which was then subjected to a Relion-4.0 3D classification. It identified a distinct subset of particles that eventually led to a 3.2 Å reconstruction (Conformation 3, similar to S_A_) with 104,849 particles using Relion-4.0 AutoRefine. The rest of the particles were then imported to CryoSPARC for 3D Classification, which led to another distinct subset of 176,488 particles being identified (Conformation 2, similar to SD), with a reconstruction of 2.9 Å using CryoSPARC homo refine. From the same 3D classification, a distinct conformation of 26S proteasome where the ATPase ring largely expanded was identified. Further 3D classification identified 3 variations of such conformation, namely S_BAG1_ (21,831 particles, 3.6 Å resolution), S_BAG2_ (16,466 particles, 3.5 Å resolution) and S_BAG3_ (19,267 particles, 3.6 Å resolution) using Non-uniform refinement in CryoSPARC.

Initial movie frame alignment and CTF estimation of the Rpn1:Bag1 complex were performed using cryoSPARC. 6,415,119 particles were picked by crYOLO using a network trained on a subset of the datasets. After multiple rounds of 2D classification, 743,645 particles were subjected to Ab-Initio Reconstruction in CryoSPARC with 5 classes. A class that looked promising was selected for further processing, which corresponded to 223,163 particles. It was subjected to Relion-5.0 focused refinement, followed by particle subtraction, focused refinement and focused classification, 2 conformations of the tenary complex of the Hsp70NBD:Bag1:Rpn1 can be identified (25,238 particles, 4.8 Å resolution; 30,256 particles, 6.2 Å).

For the control dataset 26S:BAG_UBL_ dataset, movie frame alignment was performed using cryoSPARC. For the other dataset 26S:Bag1_BAG_, movie frame alignment was performed using Relion 4.0. CTF estimation was estimated by CTFFIND4. 602,653 and 648,687 particles were initially picked by crYOLO for 26S:BAG1_UBL_ and 26S:BAG1_BAG_ datasets, respectively. After multiple rounds of 2D classification and 3D classification in Relion-4.0, followed by cryoDRGN analysis^78^, no map with expanded ATPase ring like S_BAGs_ could be found in either dataset.

### Model building

For the Bag1-bound 26S proteasome, the model was built on S_BAG1_ map. The published model (PDB ID: 7w3k) was used as initial reference. Most of the chains fit into the densities readily, whereas the ATPase chains were individually docked to the corresponding densities, followed by multiple rounds of manual adjustment in Coot ^79^ and real space refinement in PHENIX ^80^. Details of the model-building process and the quality of the final atomic models are provided in Extended Table 3.

The current model and 7w3k were aligned on the 20S for RMSD measurement of the Rpt. Per-residue RMSD measurement on the mainchain of the ATPase domain was calculated based on BioPandas (81).

## Acknowledgements

This work was funded by the Spanish State Research Agency through grants PID2022-137175NB-I00 (AEI/FEDER, UE), AEI/10.13039/501100011033), and the “Severo Ochoa” Programme for Centres of Excellence in R&D (CEX2023-001386-S) (to JMV and JC). Support was also provided by the German Research Foundation (DFG) through CRC889 (project number 154113120; project A11 to ES), and Germany’s Excellence Strategy (project number EXC 2067/1-390729940 to ES). Cryo-ET instrumentation was jointly funded by the DFG Major Research Instrumentation program (448415290) and the Ministry of Science and Culture of the State of Lower Saxony. Additional funding was provided to MM by the Centro de Investigación Biomédica en Red de Enfermedades Respiratorias (CIBERES), an initiative of the Instituto de Salud Carlos III (ISCIII). We also thank the Helmholtz Centre for Infection Research, Instruct-ERIC, and Amsterdam, Netherlands (PID 26461) for providing biomass of HEK cells. Special thanks to the European Synchrotron Radiation Facility (ESRF) in Grenoble for access to the Titan Krios cryoelectron microscope (MX2263).

## Author contribution

Protein samples were produced by MML, JM, AS and MA. MML, TCC, ES and JM performed and analysed the cryo-EM experiments. MM and MML performed and analysed the ITC experiments. JMV, JC and ES designed and supervised the research. Figures and movies were prepared by JM and ES. ES, JC and JMV wrote the manuscript with contributions from all authors.

## Data availability

Data supporting the findings of this work are available within the Article, Extended Data and the Supplementary Information files. The Cryo-EM reconstructions of the Bag1-bound 26S proteasome have been deposited in the Electron microscopy Data Bank (www.emdatabank.org) under the following accession codes; EMDB 52097 (S_BAG1_), 52193 (S_BAG2_), and 52194 (S_BAG3_). The Cryo-EM reconstructions of the ternary complex of Hsp70_NBD_:Bag1:Rpn1 are deposited under EMDB accession codes of 52195 (class 1) and 52196 (class 2). Atomic model coordinates and associated structure factors of the Bag1-bound 26S proteasome in S_BAG1_ is deposited in Protein Data Bank database (www.pdb.org) under PDB accession code 9HEU. Code used for RMSD can be accessed under (https://github.com/xtalfire/RMSD-measure)

