## Supplementary Figures for "Structures of the 26S proteasome in complex with the Hsp70 cochaperone Bag1 reveal a novel mechanism of ubiquitin-independent proteasomal degradation"

### Figs. E1 to E10

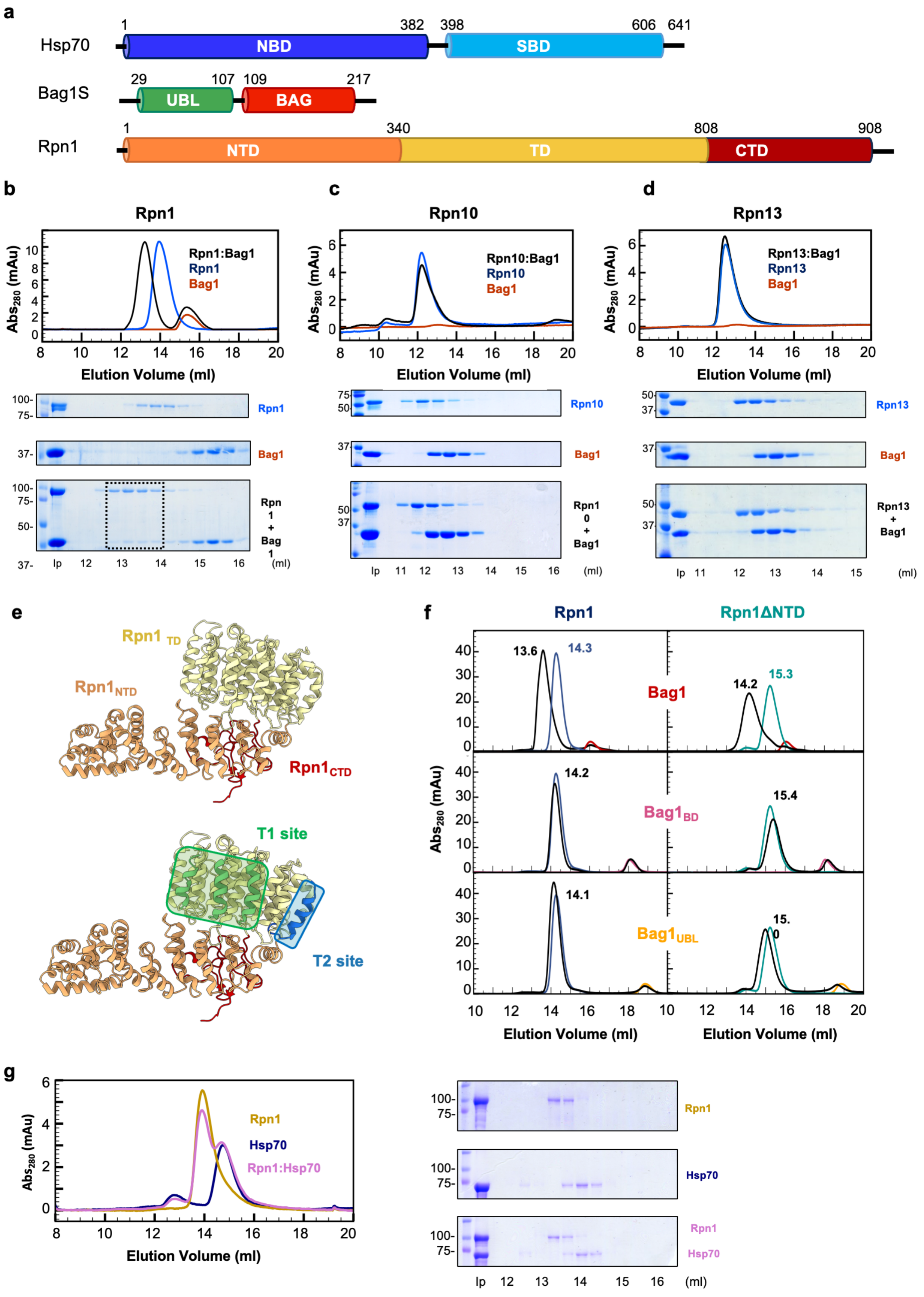

**Fig. E1. Domain structure and SEC analysis of the Bag1-Rpn1 interaction.** (**a**) Domain structures of the chaperone Hsp70, the cochaperone Bag1, and the proteasomal subunit Rpn1. **(b-d)** SEC and SDS-PAGE analysis to identify the proteasome binding subunit of Bag1. Bag1 was incubated with the proteasome ubiquitin receptors Rpn1 **(b)**, Rpn10 **(c)** and Rpn13 **(d)** and then subjected to the SEC column. Only Rpn1 **(b)** shows a peak shift in the presence of Bag1,indicating the formation of a dimeric complex. **(e)** Structure of Rpn1 highlighting its domain structures (upper panel). $Rpn1_{TD}$, $Rpn1_{NTD}$ and $Rpn1_{CTD}$ are colored in light yellow, orange and red, respectively (PDB: 7W3K). T1 ubiquitin binding site (Chen et al., 2016), and T2 UBL binding site (Shi et al., 2016) are highlighted in green and blue, respectively (lower panel). **(f)** Rpn1 interacts with the UBL domain of Bag1 ($Bag1_{UBL}$) but not the BAG domain ($Bag1_{BD}$). Deletion of the N-terminal region of Rpn1 ($Rpn1_{NTD}$) did not interfere with Bag1 binding, suggesting that $Bag1_{UBL}$ interacts with $Rpn1_{TD}$ but not $Rpn1_{NTD}$. SEC analysis to identify the domain specificity revealed a stable interaction between full-length Bag1 and Rpn1. **(g)** SEC analysis showed no peak shift when mixing Rpn1 and Hsp70, indicating no direct interaction between Rpn1 and Hsp70.

**a**

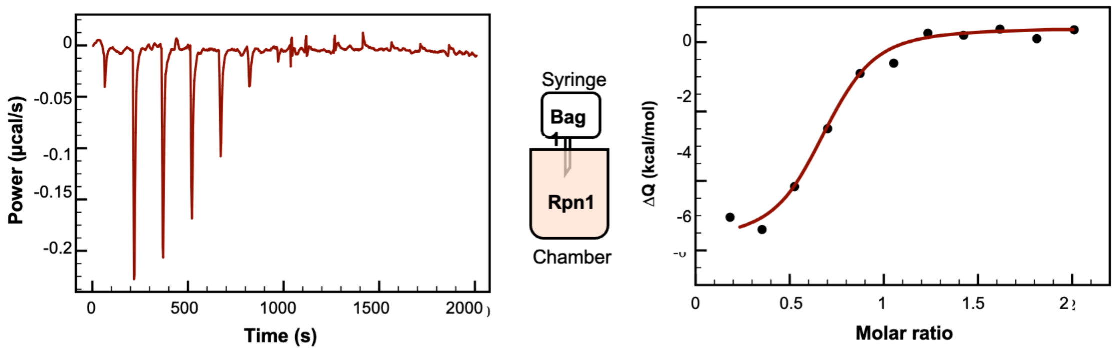


**b**

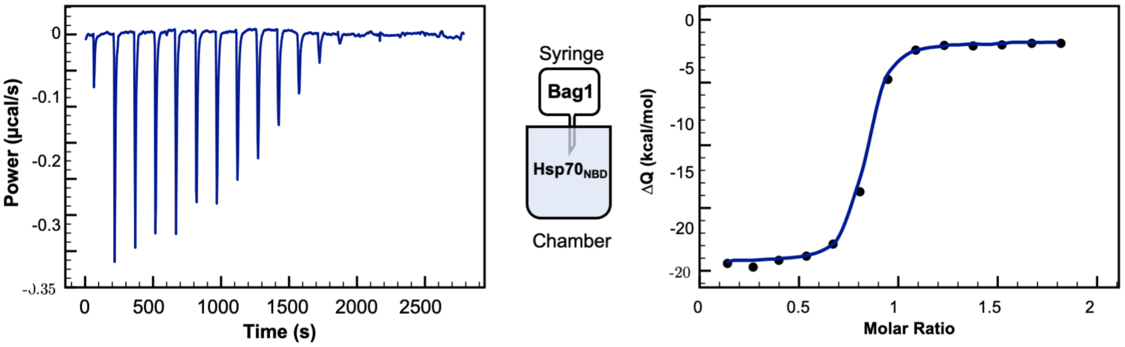


**c**

| | | |
|---|---|---|
| **Kd (M)** | (4.76 ± 1.8) ·$10^{-7}$ | (5.00 ± 1.5) ·$10^{-8}$ |
| **ΔH (kcal/mol)** | -6.8 ± 1.0 | -17.3 ± 0.1 |
| **n** | 0.59 ± 0.01 | 0.77 ± 0.01 |
| **Qdil (kcal/mol)** | 0.42 ± 0.1 | -1.9 ± 0.2 |
| **ΔG (kcal/mol)** | -8.6 ± 0.2 | -10 ± 0.2 |
| **ΔS (kcal/mol)** | 5.8 ± 4 | -25 ± 1 |

**d**

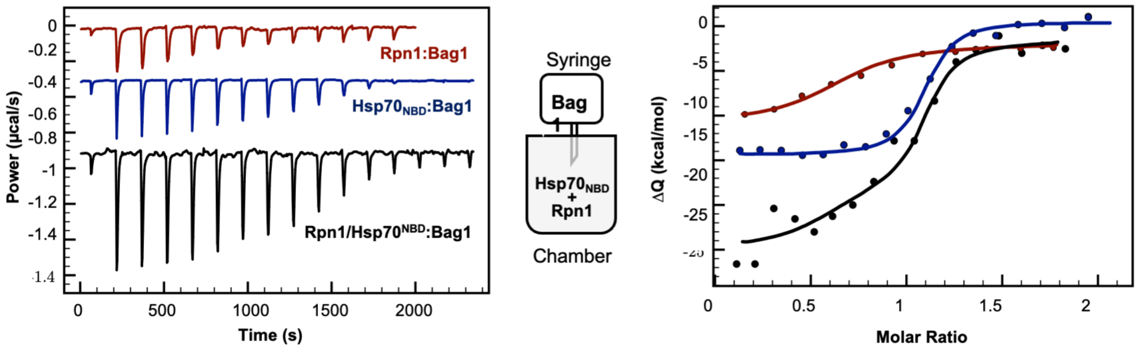


**Fig. E2. Isothermal titration calorimetry (ITC) experiment for Rpn1, Hsp70 and a mixture of Rpn1-Hsp70 titrated with Bag1.** Panels of the raw data (left) and fitted integrated data (right) of the titration experiments where Bag1 was titrated with Rpn1 **(a)**, $Hsp70_{NBD}$ **(b)**. An experimental schematic is shown between panels. All ITC binding assays were performed with an PEAQ-ITC microcalorimeter (Malvern) at 25 °C in a buffer containing 20 mM phosphate buffer pH 7.4 and 150 mM KCl, and data were analyzed using the AFFINImeter software. **(c)** Summary of the thermodynamic parameters of Rpn1:Bag1 (salmon column on the left) and $Hsp70_{NBD}$:Bag1 (blue column on the right), derived from the one-to-one binding model. Averages over multiple measurements are listed. **(d)** ITC assay for the ternary complex with Bag1 titrated against a mixture of Rpn1 and $Hsp70_{NBD}$. The curve was fitted simultaneously with binary complexes assuming independent binding of $Hsp70_{NBD}$ and Rpn1 to Bag1. The black trace in **(d)** represents the simulated formation of the ternary complex of $Hsp70_{NBD}$:Bag1:Rpn1, compared to the binding parameters in **(c)**. The individual plots for Bag1 titrated with Rpn1 and Hsp70 are also shown in red and blue, respectively.

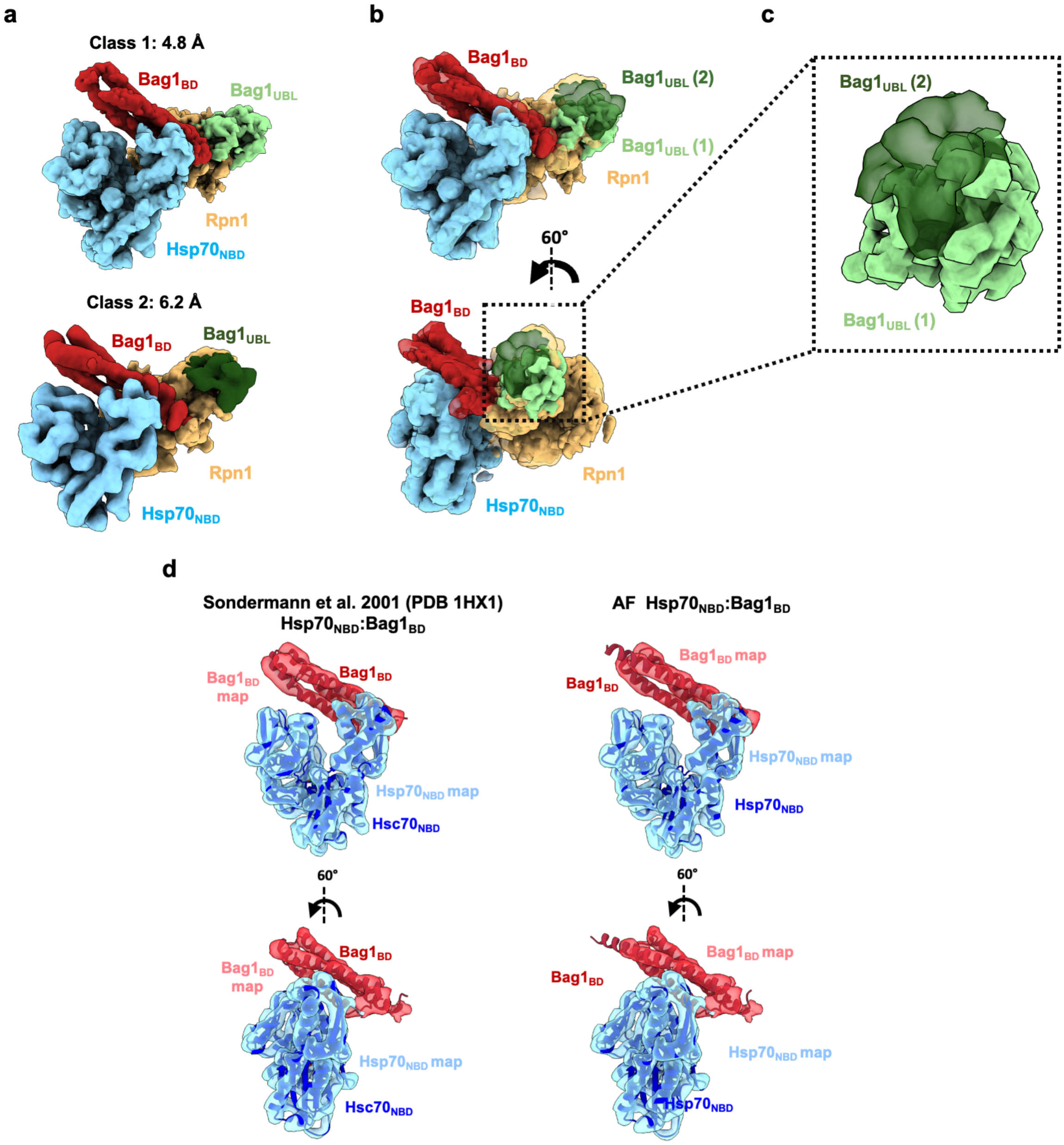


**Fig. E3. 3D reconstructions of the $Hsp70_{NBD}$:Bag1:Rpn1 ternary complex. (a)** Cryo-EM density maps of the $Hsp70_{NBD}$:Bag1:Rpn1 ternary complex (Class 1 EMDB: 52195; Class 2 EMDB: 52196). Rpn1, $Bag1_{BD}$ and $Hsp70_{NBD}$ are colored in beige, red and blue, respectively in both maps. The $Bag1_{UBL}$ from Class 1 is depicted in light green and in dark green for Class 2. **(b)** An overlay of the two maps reveals their structural similarity, with the exception of the $Bag1_{UBL}$ domain, which displays structural flexibility **(c)**. **(d)** Comparison of the $Hsp70_{NBD}$:$Bag1_{BD}$ interaction (blue:red) in the context of the ternary complex. Crystal structure of $Hsc70_{NBD}$:$Bag1_{BD}$ (PDB: 1HX1, Sondermann *et al*, 2001) (left) and an AlphaFold prediction, showing no noticeable differences (right). All structures were docked into the cryo-EM map of the $Hsp70_{NBD}$:Bag1:Rpn1 ternary complex.

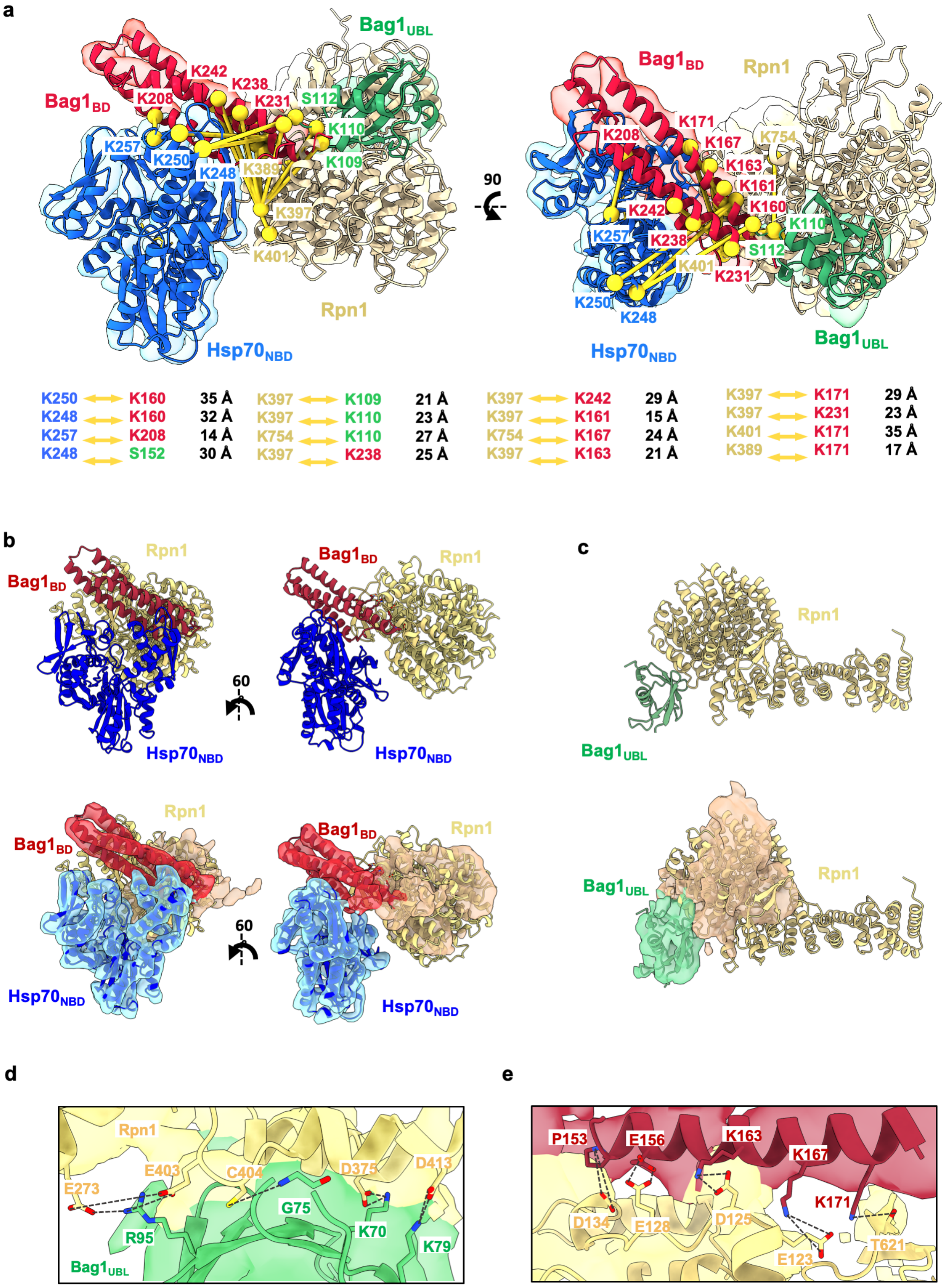

a
Bag1$_{BD}$
Bag1$_{UBL}$
Rpn1
Hsp70$_{NBD}$
K242
K238
K231
K208
S112
K110
K109
K257
K250
K248
K389
K397
K401
K171
K167
K163
K161
K160
K754
90
K250 ⟷ K160 35 Å
K248 ⟷ K160 32 Å
K257 ⟷ K208 14 Å
K248 ⟷ S152 30 Å
K397 ⟷ K109 21 Å
K397 ⟷ K110 23 Å
K754 ⟷ K110 27 Å
K397 ⟷ K238 25 Å
K397 ⟷ K242 29 Å
K397 ⟷ K161 15 Å
K754 ⟷ K167 24 Å
K397 ⟷ K163 21 Å
K397 ⟷ K171 29 Å
K397 ⟷ K231 23 Å
K401 ⟷ K171 35 Å
K389 ⟷ K171 17 Å
b
60
c
d
E273
E403
C404
D375
D413
G75
R95
K70
K79
e
P153
E156
K163
K167
K171
D134
E128
D125
E123
T621

**Fig. E4. Structural validation of the $Hsp70_{NBD}$:Bag1:Rpn1 ternary complex**. **(a)** Two views of the ternary complex reconstruction incorporating cross-links (yellow) identified through XL-MS analysis. The distances of all cross-links calculated from the structural model (rigid body fitting of atomic model of PDB: 1HX1) are also indicated. **(b)** AlphaFold prediction of Rpn1:$Bag1_{BD}$:$Hsp70_{NBD}$ complex (upper panel) fitted into our cryo-EM reconstruction of the ternary complex (EMDB: 52195) (lower panel). **(c)** AlphaFold prediction of the Rpn1:$Bag1_{UBL}$ complex (upper panel) fitted into our cryo-EM reconstruction of the ternary complex (EMDB: 52195) (lower panel). AlphaFold predicted contacts between Rpn1:$Bag1_{UBL}$ **(c)** and Rpn1:$Bag1_{BD}$ **(d).** Putative binding interfaces extracted from the predictions stated above. Color code follows Rpn1 in beige/yellow, $Bag1_{UBL}$ in green, $Bag1_{BD}$ in red and $Hsp70_{NBD}$ in blue.

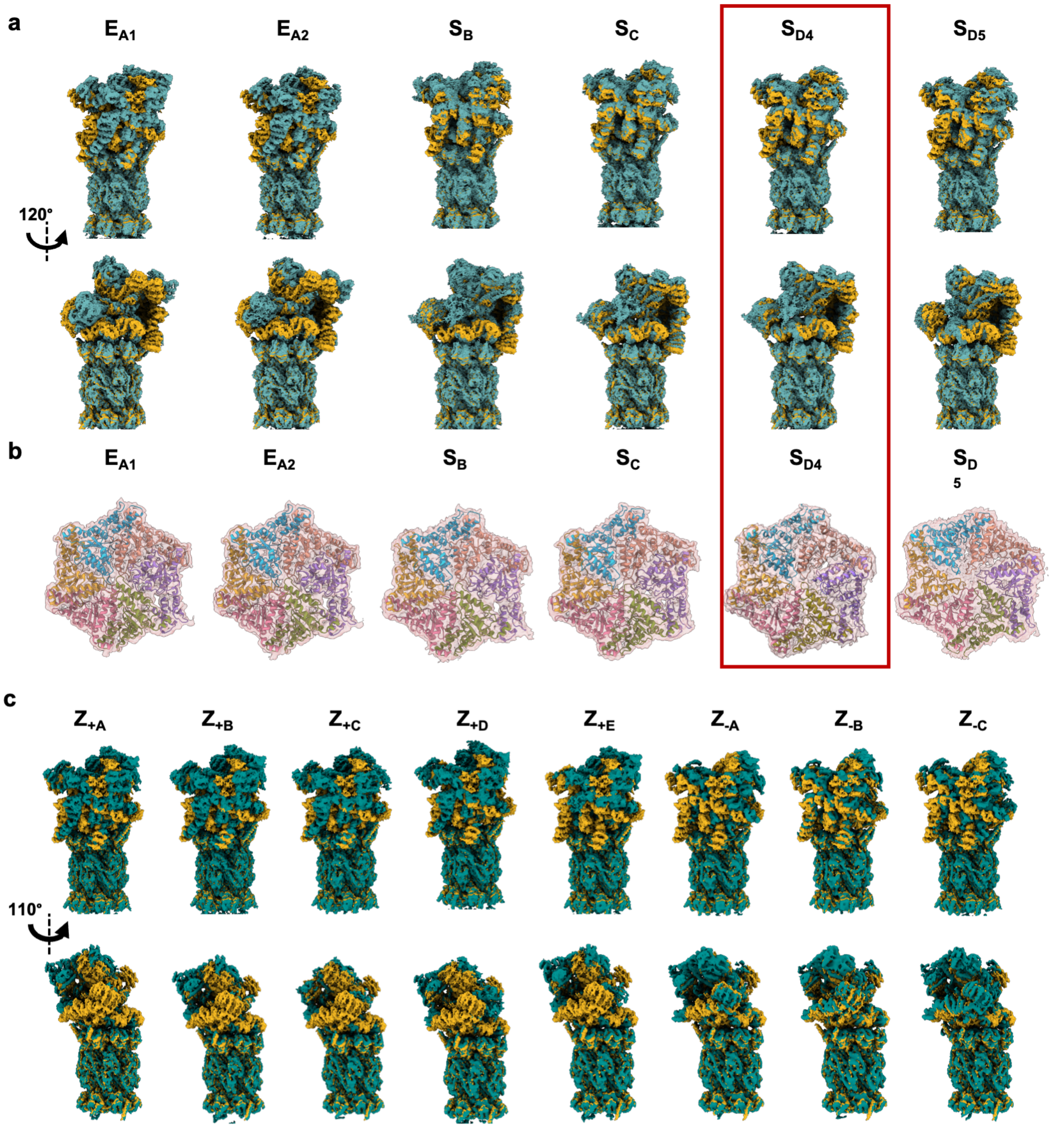


**Fig. E5. Structural comparison of the Bag1-bound 26S proteasome with reported human 26S proteasome structures. (a)** Structural comparison of the cryo-EM reconstruction of 26S proteasome in $S_{BAG1}$ state (goldenrod) with previously reported cryo-EM reconstructions of USP14-bound 26S proteasome (teal). Cryo-EM maps from Zhang et al. (Zhang, Zou et al. 2022) were used for comparison: $E_{A1}$ (EMDB: 32272), $E_{A2}$ (emdb: 32273), $S_B$ (EMDB: 32281), $S_C$ (EMDB: 32282), $S_{D4}$ (EMDB: 32283), $S_{D5}$ (EMDB: 32284). **(b)** Cryo-EM segmentation of the ATPase ring in previously reported structures of the USP14-bound 26S proteasome. Different from the Bag1-bound 26S proteasome structures, the tight packing of the ATPase rings was observed in all canonical structures. **(c)** Structural comparison of the cryo-EM reconstruction of 26S proteasome in $S_{BAG1}$ state (goldenrod) with previously reported cryo-EM reconstructions of the ZFAND5-bound 26S proteasome (teal). Cryo-EM maps from Lee et al. (Lee et al. 2023) were used for comparison: $Z_{+A}$ (EMDB: 14201), $Z_{+B}$ (EMDB: 14202), $Z_{+C}$ (EMDB: 14203) , $Z_{+D}$ (EMDB: 14204), $Z_{+E}$ (EMDB: 14205), $Z_{-A}$ (EMDB: 14209), $Z_{-B}$ (EMDB: 14210), $Z_{-C}$ (EMDB: 14212).

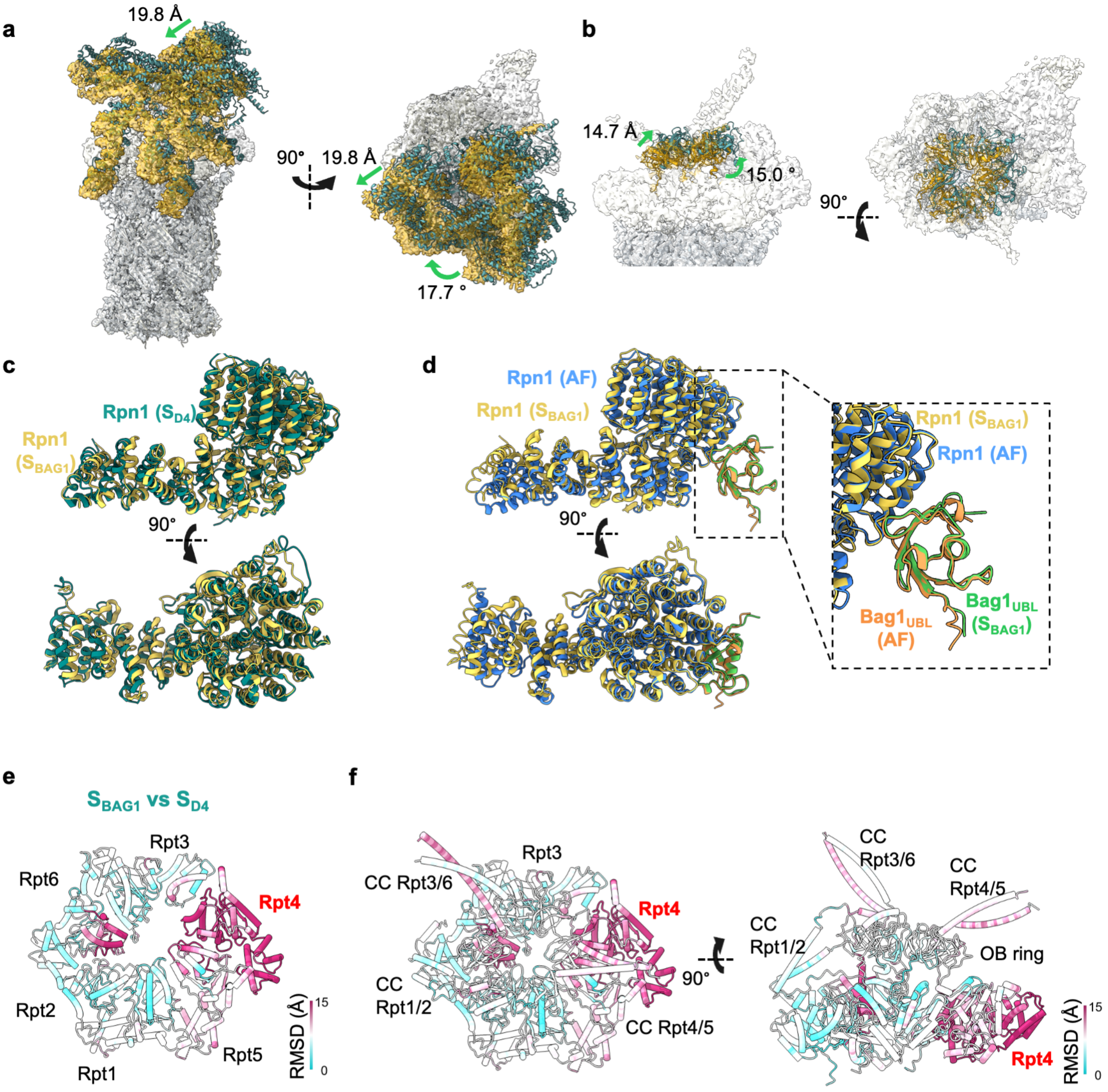


**Fig. E6. Structural comparison of the cryo-EM reconstruction of the 26S proteasome in $S_{BAG1}$ state with $S_{D4}$ state.** Structural comparison of the cryo-EM reconstruction of the 26S proteasome in $S_{BAG1}$ (EMDB: 52097 in goldenrod) with the $S_{D4}$ state (PDB: 7W3K in teal), focusing on the lid subcomplex **(a)** and OB ring **(b)**. The rest of densities are shown in light grey. The Changes in shift (Å) and angle (°) are indicated. **(c)** Structural comparison of Rpn1 in $S_{BAG1}$ (goldenrod) and $S_{D4}$ (blue green) states. No significant conformational change was observed in Rpn1. **(d)** Structural comparison of the Rpn1 at the $Bag1_{UBL}$ interface in $S_{BAG1}$ (goldenrod) with AlphaFold prediction (blue). $Bag1_{UBL}$ is shown in green ($S_{BAG1}$) and orange ($S_{D4}$), respectively. **(e,f)** Residue-wise RMSD of the Rpt subunits calculated from the main chain including N, Cα, C, O between $S_{BAG1}$ and $S_{D4}$ states. Residue-wise RMSDs of the main-chain of the ATPase domain **(e)** and whole Rpt subunits **(f),** compared the difference from $S_{BAG1}$ with $S_{D4}$ (PDB: 7W3K), illustrates the extent of the conformational shift caused by Bag1-binding (in Å). Two structures are aligned to the α ring of the CP. The RMSD is plotted on the $S_{BAG1}$ structure. Rpt4 (large, small domains), Rpt5 (large), Rpt3 (small) and Rpt6 (large) shows large shifts in the ATPase ring. The coiled-coil (CC) of the Rpt3/6 and Rpt4/5 are displaced compared to the Rpt1/2 CC.

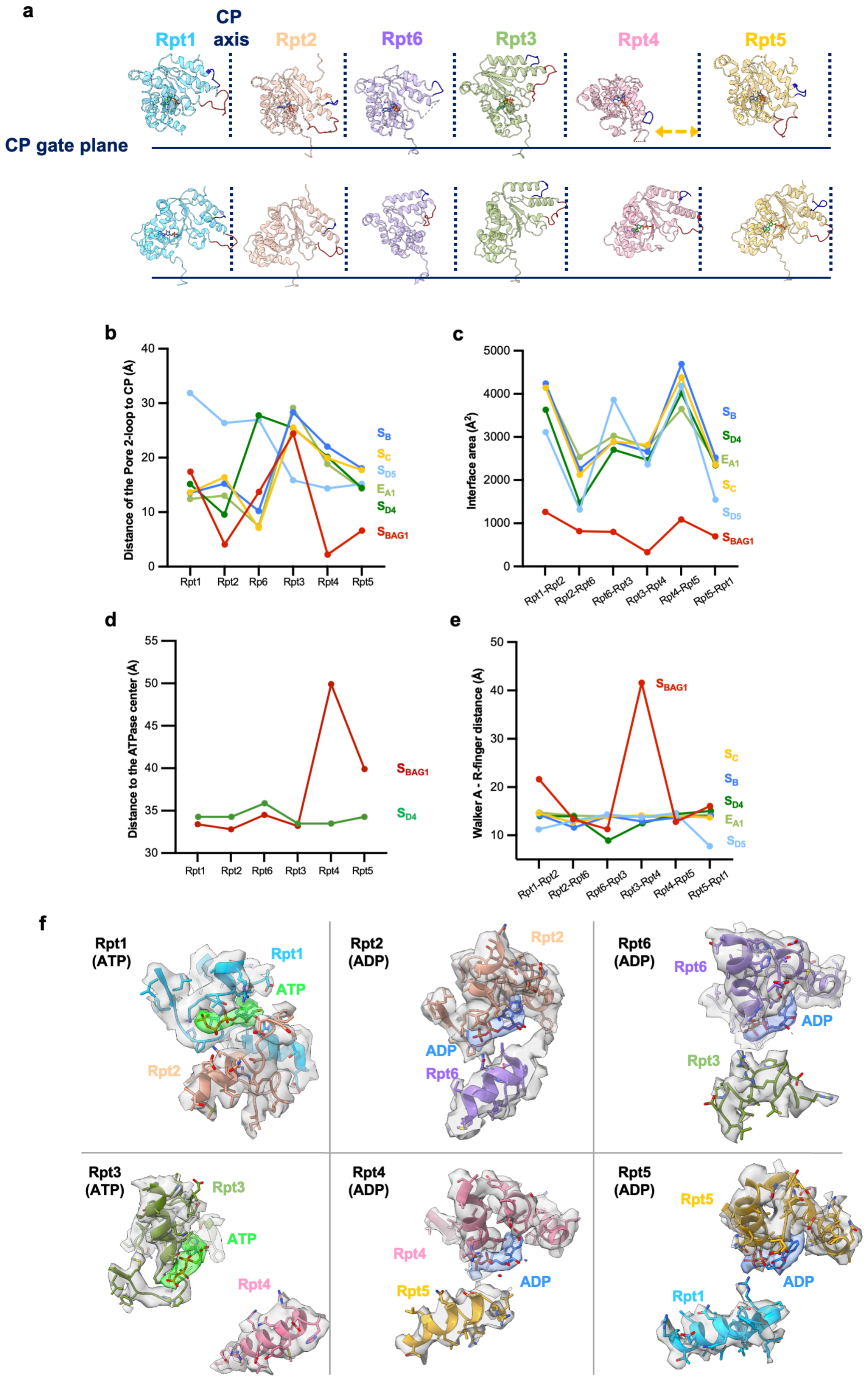
a
CP axis
Rpt1
Rpt2
Rpt6
Rpt3
Rpt4
Rpt5
CP gate plane
b
Distance of the Pore 2-loop to CP (Å)
Rpt1 Rpt2 Rp6 Rpt3 Rpt4 Rpt5
$S_B$
$S_C$
$S_{D5}$
$E_{A1}$
$S_{D4}$
$S_{BAG1}$
c
Interface area (Å²)
Rpt1-Rpt2 Rpt2-Rpt6 Rpt6-Rpt3 Rpt3-Rpt4 Rpt4-Rpt5 Rpt5-Rpt1
d
Distance to the ATPase center (Å)
e
Walker A - R-finger distance (Å)
f
Rpt1 (ATP)
Rpt2 (ADP)
Rpt6 (ADP)
Rpt3 (ATP)
Rpt4 (ADP)
Rpt5 (ADP)
ATP
ADP

**Fig. E7. Structural analysis of the ATPase Rpt subunits in $S_{BAG1}$. (a)** Organization of the AAA+ ATPase and interaction with the CP. Individual Rpt subunits in $S_{BAG1}$ (upper panel, PDB: 9HEU) and $S_{D4}$ (lower panel, PDB: 7W3K) rotated around the CP axis are shown in approximately 60° steps. Due to the deformation of the ring structure, orientation of each subunit in $S_{BAG1}$ is adjusted to properly show the side view and pore-2 loop. The positions of the CP axis and CP plane of the gate are shown as dotted lines and solid lines, respectively. Pore-1 loops and Pore-2 loops of each subunit are highlighted in blue and red, respectively. The Bag1-induced conformational change alters the position of Rpt4 subunit far away from the central axis. **(b)** Plot of distances between the closest α carbon of the Pore-2 loops to the plane of the CP gate. Pore-2 loops in Rpt2, Rpt4 and Rpt5 in $S_{BAG1}$ are close to the CP plane compared to any of reported structures($E_{A1}$, $S_B$, $S_C$, $S_{D4}$, $S_{D5}$) (Zhang, Zou et al. 2022). **(c)** Contact area between the large AAA+ domains of neighboring Rpt subunits in $S_{BAG1}$ and the reported states ($E_{A1}$, $S_B$, $S_C$, $S_{D4}$, $S_{D5}$) (Zhang, Zou et al. 2022). All interfaces in $S_{BAG1}$ shows small contact interface. **(d)** Distance of the centroid of individual subunits to the axis of the ATPase ring in $S_{BAG1}$ and $S_{D4}$. Rpt4 is completely off the axis as it is shown in Figure 3g. **(e)** Opening of the nucleotide-binding pockets in $S_{BAG1}$ and the reported states ($E_{A1}$, $S_B$, $S_C$, $S_{D4}$, $S_{D5}$) (Zhang, Zou et al. 2022). The distances between the α carbon of Walker-A Thr residue and the α carbon of the Arg finger of the neighboring subunit are shown. The interface between Rpt3 and Rpt4 in $S_{BAG1}$ is especially opened. **(f)** Detailed views of individual Rpt nucleotide-binding pockets, together with the adjacent Rpt subunit. Nucleotide binding pocket of a subunit and the closest α helix of the neighboring subunit are shown. In Rpt1 and Rpt3 pockets, ATP densities are observed, while ADP was observed in Rpt2, 4, 5 and 6.

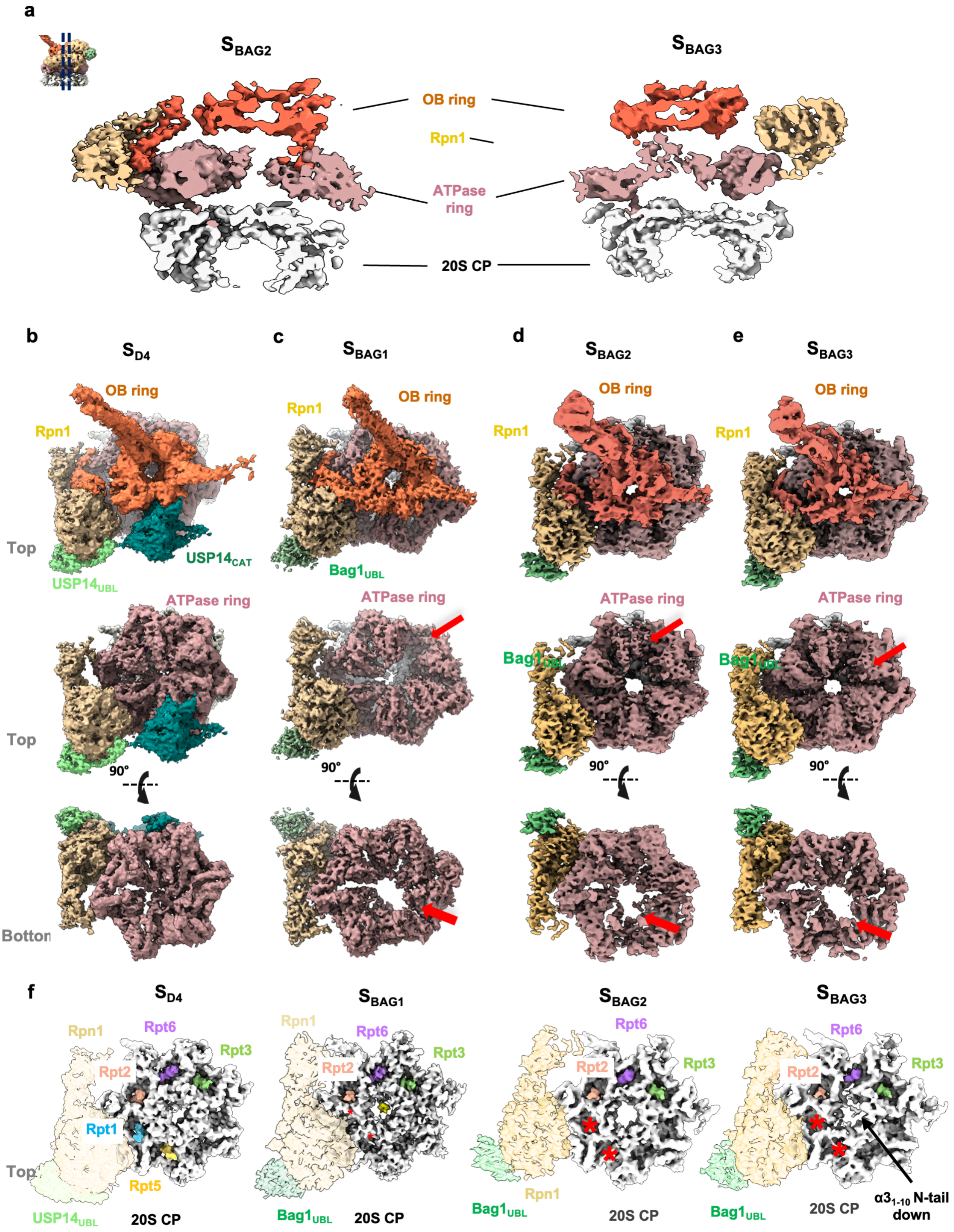

a
S_BAG2
S_BAG3
OB ring
Rpn1
ATPase ring
20S CP
b
S_D4
c
S_BAG1
d
S_BAG2
e
S_BAG3
OB ring
Rpn1
Top
USP14_CAT
USP14_UBL
Bag1_UBL
ATPase ring
Bag1_UBL
90°
Bottom
f
S_D4
S_BAG1
S_BAG2
S_BAG3
Rpt6
Rpt3
Rpt2
Rpt1
Rpt5
USP14_UBL
20S CP
Bag1_UBL
α3_1-10 N-tail down

**Fig. E8. Cryo-EM segmentations of the 26S proteasome in $S_{D4}$ and $S_{BAG1-3}$. (a)** Cross-sections of cryo-EM maps in $S_{BAG2}$ (left panel) and $S_{BAG3}$ (right panel) in the same orientation and color with Figure 4a, focusing on the interface between the ATPase and CP rings. The color codes are as follows; OB ring (orange), ATPase ring (rosy brown), 20S (white) and Rpn1 (beige). In both $S_{BAG2}$ and $S_{BAG3}$ structures, a significant cavity is created in the ATPase domain as it is observed in $S_{BAG1}$. **(b-e)** Structural comparison of the cut-away views of the USP14-bound 26S proteasome ($S_{D4}$) (EMDB: 32282) **(b),** Bag1-bound 26S proteasome in $S_{BAG1}$ (EMDB: 52097) **(c)**, Bag1-bound 26S proteasome in $S_{BAG2}$ (EMDB: 52193) **(d)** and in $S_{BAG3}$ (EMDB: 52194) **(e)**. The same color codes are used as in panel **(a)**. Each segment is shown down the long axis of the 26S proteasome from the OB ring (top) and the ATPase domain ring (top view; middle and bottom view; bottom). Large cavities are observed at the center of the ATPase ring in all three $S_{BAG}$ conformations. **(f)** Insertion of Rpt C-terminal tails into CP α ring pockets. The top views of the 20S CP, Rpn1, as well as $Bag1_{UBL}$ domains are shown in white, tan, and light green, respectively. The C-terminal Rpt densities inserted into the CP are colored in Rpt1 in deep sky blue, Rpt2 in salmon pink, Rpt6 in purple, Rpt3 in olive green, and Rpt5 in goldenrod. Only three C-terminal Rpt tails (Rpt2, Rpt3, Rpt5) are inserted into CP α pockets in $S_{BAG}$ conformations.

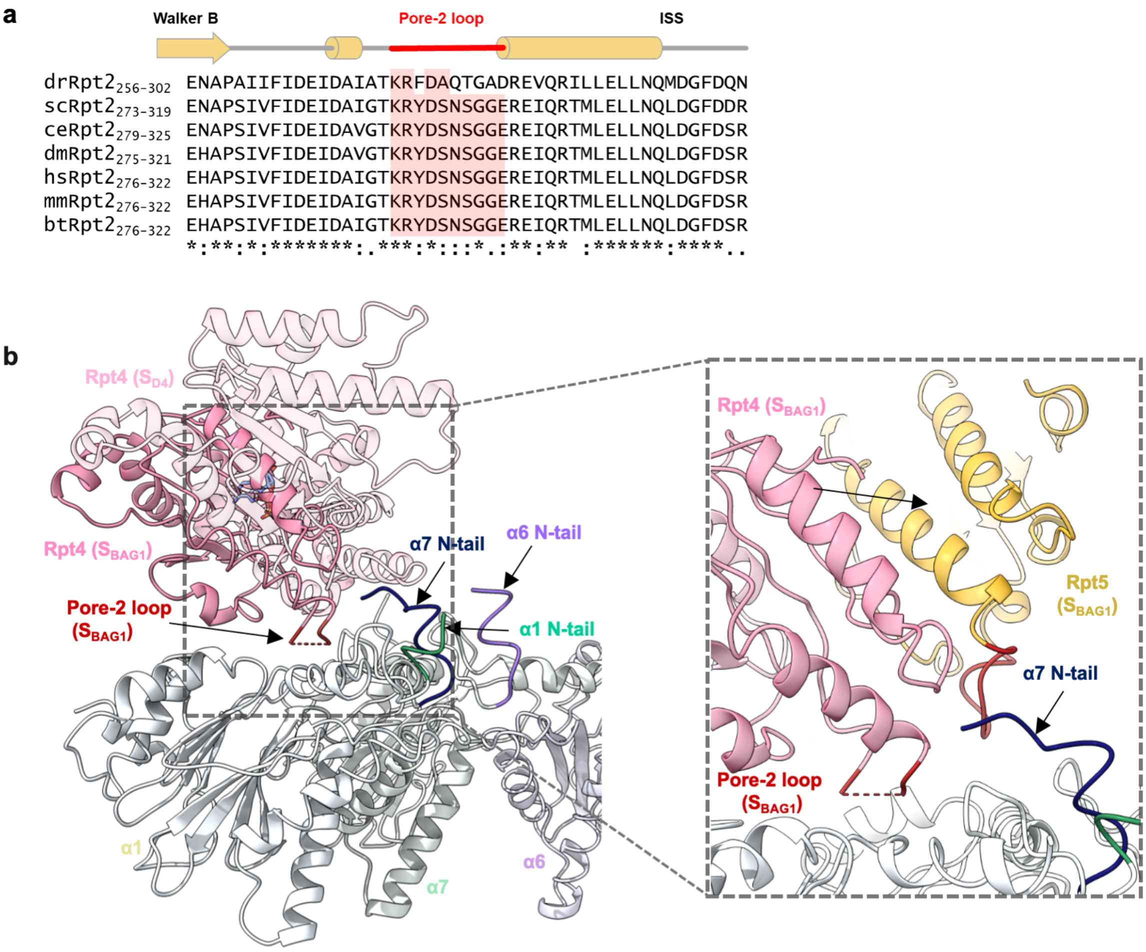


**Fig. E9. Conserved sequence of Rpt2 Pore-2 loop and structural change of the Rpt4 ATPase domain.** **(a)** Sequence alignment of Rpt2 subunits across different species showing conserved sequence. The Rpt2 Pore-2 loop is highly conserved across species except drosophila. **(b)** Large movement of Rpt4 brings the Pore-2 loop above the α pockets between α6 and α7, although the 241-249 residues density is not observed due to the flexibility. However, the N-terminal segments of α6 and α7 are kept in the up position. Detailed view is shown in the dotted square.

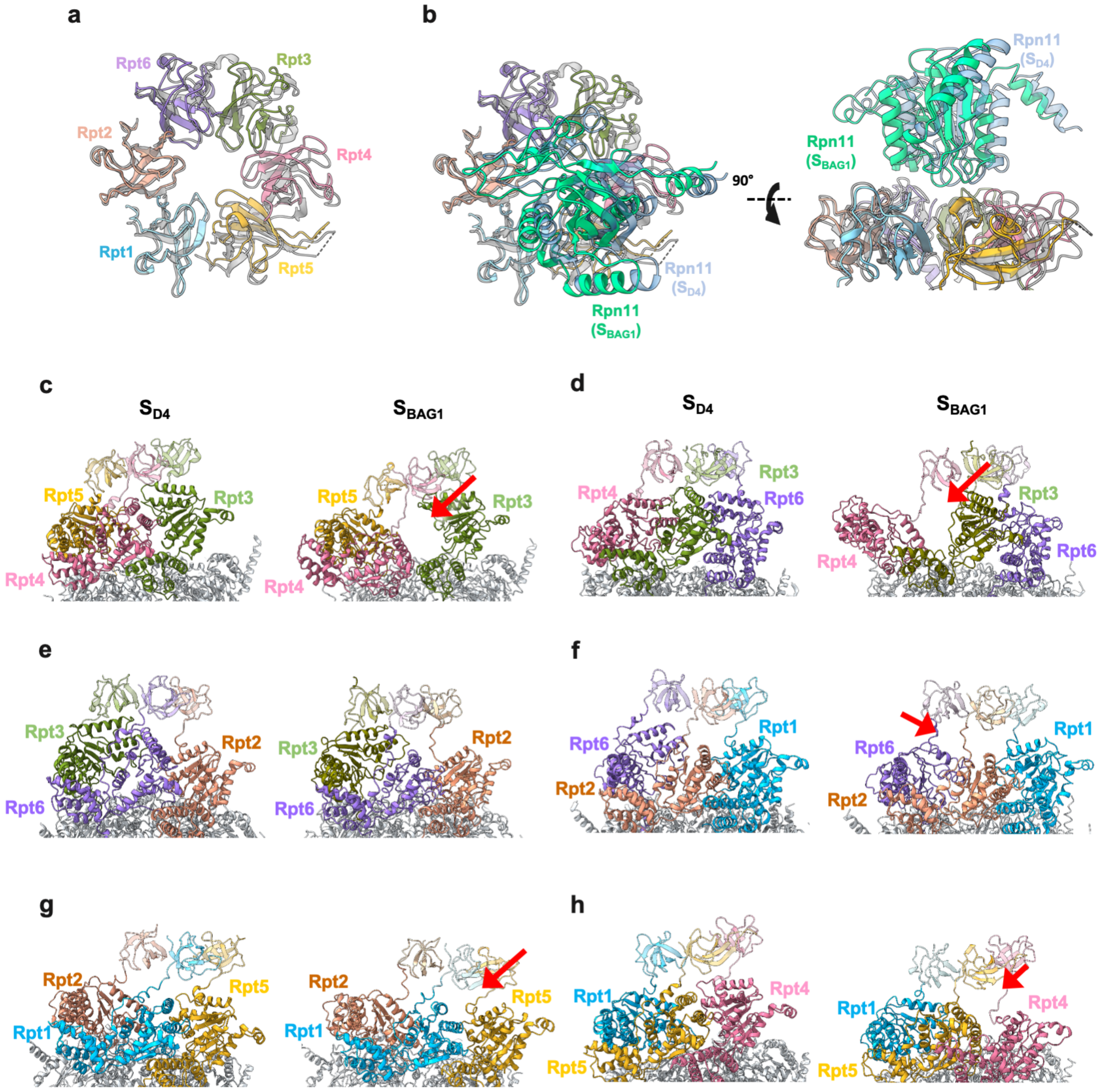


**Fig. E10. The Interfaces between OB and ATPase rings in $S_{BAG1}$ and $S_{D4}$ are compared.** Structural compari2sons between $S_{D4}$ (PDB: 7W3K) and $S_{BAG1}$ (PDB: 9HEU) and, focusing on the OB domains **(a)** and Rpn11 **(b)**. No significant changes between both states are observed in the OB ring. Rpt subunits of $S_{BAG1}$ are colored with Rpn1 in deep sky blue, Rpt2 in salmon pink, Rpt6 in purple, Rpt3 in olive green, Rpt4 in pink and Rpt5 in goldenrod, whereas $S_{D4}$ subunits are in gray. Rpn11 is depicted in turquoise for $S_{BAG1}$ and light blue for $S_{D4}$. **(c-h)** Comparison of the OB-ATPase interfaces formed by three sequential Rpt subunits. The interfaces are shown in every 60° step. Rpt1, Rpt2, Rpt6, Rpt3, Rpt4 and Rpt5 are shown in deep sky blue, salmon, purple, olive green, pink and goldenrod. 20S CP is shown in white. The OB domains are shown in transparent. Opened interfaces of the OB-ATPase ring are highlighted by red arrows. Due to the significant conformational change of the ATPase domains, a large opening is created at the interface between OB and ATPase rings.
